# Genetic signatures of human cytomegalovirus variants acquired by seronegative glycoprotein B vaccinees

**DOI:** 10.1101/432922

**Authors:** Cody S. Nelson, Diana Vera Cruz, Melody Su, Guanhua Xie, Nathan Vandergrift, Robert F. Pass, Michael Forman, Marie Diener-West, Katia Koelle, Ravit Arav-Boger, Sallie R. Permar

## Abstract

Human cytomegalovirus (HCMV) is the most common congenital infection worldwide, and a frequent cause of hearing loss or debilitating neurologic disease in newborn infants. Thus, a vaccine to prevent HCMV-associated congenital disease is a public health priority. One potential strategy is vaccination of women of child-bearing age to prevent maternal HCMV acquisition during pregnancy. The glycoprotein B (gB) + MF59 adjuvant subunit vaccine is the most efficacious tested clinically to date, demonstrating approximately 50% protection against HCMV infection of seronegative women in multiple phase 2 trials. Yet, the impact of gB/MF59-elicited immune responses on the population of viruses acquired by trial participants has not been assessed. In this analysis, we employed quantitative PCR as well as multiple sequencing methodologies to interrogate the magnitude and genetic composition of HCMV populations infecting gB/MF59 vaccinees and placebo recipients. We identified several differences between the viral dynamics of acutely-infected vaccinees and placebo recipients. First, there was reduced magnitude viral shedding in the saliva of gB vaccinees. Additionally, employing a panel of tests for genetic compartmentalization, we noted tissue-specific gB haplotypes in the majority of vaccinees though only in a single placebo recipient. Finally, we observed reduced acquisition of genetically-related gB1, gB2, and gB4 genotype “supergroup” HCMV variants among vaccine recipients, suggesting that the gB1 genotype vaccine construct may have elicited partial protection against HCMV viruses with antigenically-similar gB sequences. These findings indicate that gB immunization may have had a measurable impact on viral intrahost population dynamics and support future analysis of a larger cohort.

**Author Summary:** Though not a household name like Zika virus, human cytomegalovirus (HCMV) causes permanent neurologic disability in one newborn child every hour in the United States - more than Down syndrome, fetal alcohol syndrome, and neural tube defects combined. There are currently no established effective preventative measures to inhibit congenital HCMV transmission following acute or chronic HCMV infection of a pregnant mother. However, the glycoprotein B (gB) vaccine is the most effective HCMV vaccine tried clinically to date. Here, we utilized high-throughput, next-generation sequencing of viral DNA isolated from patients enrolled in a gB vaccine trial, and identified several impacts that this vaccine had on the size, distribution, and composition of the *in vivo* viral population. These results have increased our understanding of why the gB/MF59 vaccine was partially efficacious and will inform future rational design of a vaccine to prevent congenital HCMV.

## Introduction

Human cytomegalovirus (HCMV) congenital infection affects 1 in 150 pregnancies (1) and is the most frequent non-genetic cause of sensorineural hearing loss and neurodevelopmental delay in infants worldwide (2). Additionally, HCMV is the most common infectious agent among allograft recipients, often causing end-organ disease such as hepatitis, pneumonitis, or gastroenteritis and predisposing the individual to graft rejection (3). It has been estimated that an efficacious HCMV vaccine would save the United States 4 billion dollars and 70,000 quality-adjusted life years annually, and thus HCMV vaccine development has remained a tier 1 priority of the National Academy of Medicine for the past 17 years (4).

The glycoprotein B (gB) + MF59 adjuvant vaccine is the most efficacious HCMV vaccine platform trialed to date, demonstrating partial vaccine protection in multiple patient populations. In a cohort of HCMV-seronegative postpartum women, gB vaccination achieved a promising 50% vaccine efficacy (5). When this study was subsequently repeated in a cohort of seronegative adolescent women, a comparable level of vaccine-protection was observed (6). Furthermore, in allograft recipients, the same gB vaccine reduced duration of HCMV viremia and antiviral therapy (7). The mechanism of this partial vaccine protection remains unknown, though we and others have observed this vaccine platform was particularly poor at eliciting heterologous neutralizing antibodies in these populations and that non-neutralizing antibody responses may have played a role (8, 9). Previously, we reported population-level virus sequencing of HCMV hypervariable genes in this cohort, revealing no differences in virus phylogeny between vaccinees and placebo recipients (10). However, in light of recent discoveries regarding the high intrahost diversity of *in vivo* HCMV populations (11–13), a critical question in the field remains whether vaccination exerts immune pressure on viral populations resulting in any distinction between viruses acquired by gB/MF59 vaccinees and placebo recipients.

With a genome consisting of 236 kilobase pairs (14) and encoding approximately 164 genes (15), HCMV has the largest genome of any human virus. It is well established that HCMV is highly polymorphic between and within individuals, defined via a variety of sequencing methodologies including restriction fragment length polymorphism analysis (16), targeted gene sequencing (10, 17–21), and whole genome sequencing (11 -13). Yet the source of this diversity remains poorly understood. If multiple unique viral variants are identified in a single individual (so-called “mixed infection”), does this represent *de novo* mutations, simultaneous initial infection with multiple unique variants, or independent, sequential infection events? Mixed infections have been frequently detected in both chronically HCMV-infected individuals (22) and immunocompromised hosts (18–20). Yet, we previously described that recently-seroconverted women from the gB/MF59 vaccine trial predominantly had a single virus strain detected in all tissues and at time points up to 3.5 years (10) when evaluated by a traditional Sanger sequencing methodology, suggesting that mixed infections in healthy individuals may result from independent, sequential infection events. Of note, the pathologic relevance of mixed infection (detected by PCR) is purely theoretical, as the minor variant in a mixed population of viruses has never been successfully isolated in tissue culture.

One major limitation of traditional HCMV genotyping to quantify HCMV diversity is a lack of sensitivity to detect viral variants present at low frequency. Recent HCMV whole-genome, next-generation sequencing (NGS) has suggested that there is remarkable intrahost diversity, stemming from the presence of low-frequency alleles representing minor viral variants (11–13). Thus, it has been established that HCMV likely exists within individual hosts as a heterogeneous population of related viral variants (13). Subsequent characterization of the intrahost composition and distribution of low-frequency viral variants, has led to recognition of unique viral populations between individual organs representing anatomic compartmentalization of viral populations (12).

As HCMV diversity is due to the presence of distinct, low-frequency viral variants, traditional sequencing methodologies may not be the most sensitive means to discern differences between intrahost viral populations. Thus, we applied a previously-validated (23) sequencing methodology and analysis pipeline termed Short NGS Amplicon Population Profiling (SNAPP) to investigate the viral populations of recently-seroconverted gB/MF59 vaccinees and placebo recipients. This technique, which employs sequencing of an approximately 500 base-pair region at tremendous read depth, has facilitated a more complete understanding of *in vivo* viral dynamics. We hypothesize that gB/MF59 vaccination limited the complexity of the viral population following primary HCMV infection. The study of HCMV population dynamics including viral load, pairwise genetic diversity, number of unique haplotypes (viral variants), and the characteristics of those variants may yield a more comprehensive understanding of the mechanism of partial vaccine efficacy.

## Results

### Viral load, number of haplotypes, and sequence diversity by vaccine group

We obtained HCMV DNA extracted from whole blood, saliva, urine, and vaginal fluid of 32 gB/MF59 vaccinee and placebo recipients following HCMV primary infection. Samples were taken approximately monthly, though the sampling timeline was heterogeneous between trial participants. Considering data from all anatomic compartments, viral load ranged from undetectable to 4.7 log_10_ DNA copies/mL. Around the time of seroconversion, mean viral load in vaccine recipients was 0.85 times that in placebo recipients (Table 1), although this difference was not statistically significant in a regression analysis that accounted for multiple samples per patient (95% CI: 0.23 times to 3.07 times, p=0.792).

**Table 1.**
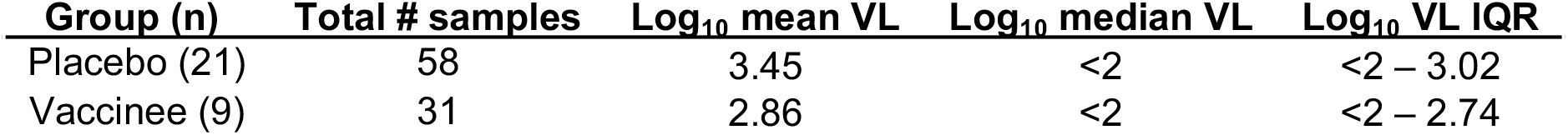
Viral load distribution (log_10_ copies/ml) by vaccination status at time of seroconversion.

| **Group (n)** | **Total # samples** | **$\log_{10}$ mean VL** | **$\log_{10}$ median VL** | **$\log_{10}$ VL IQR** |
| --- | --- | --- | --- | --- |
| Placebo (21) | 58 | 3.45 | <2 | <2 – 3.02 |
| Vaccinee (9) | 31 | 2.86 | <2 | <2 – 2.74 |

Peak viremia (Figure 1A) and peak viral shedding in saliva, urine, and vaginal fluid (Figure 1B-D) was identified for each patient and separated by vaccine group. The peak levels of viremia following primary HCMV acquisition were not significantly different between vaccinee and placebo recipients. Peak urine and vaginal fluid shedding were also not significantly different between placebo and gB/MF59 vaccinees. However, the median peak saliva shedding was noted to be 6.24 times reduced in gB vaccine recipients compared to placebo (p=0.022, Friedman test + post hoc Exact Wilcoxon Rank Sum test).

**Figure 1.**
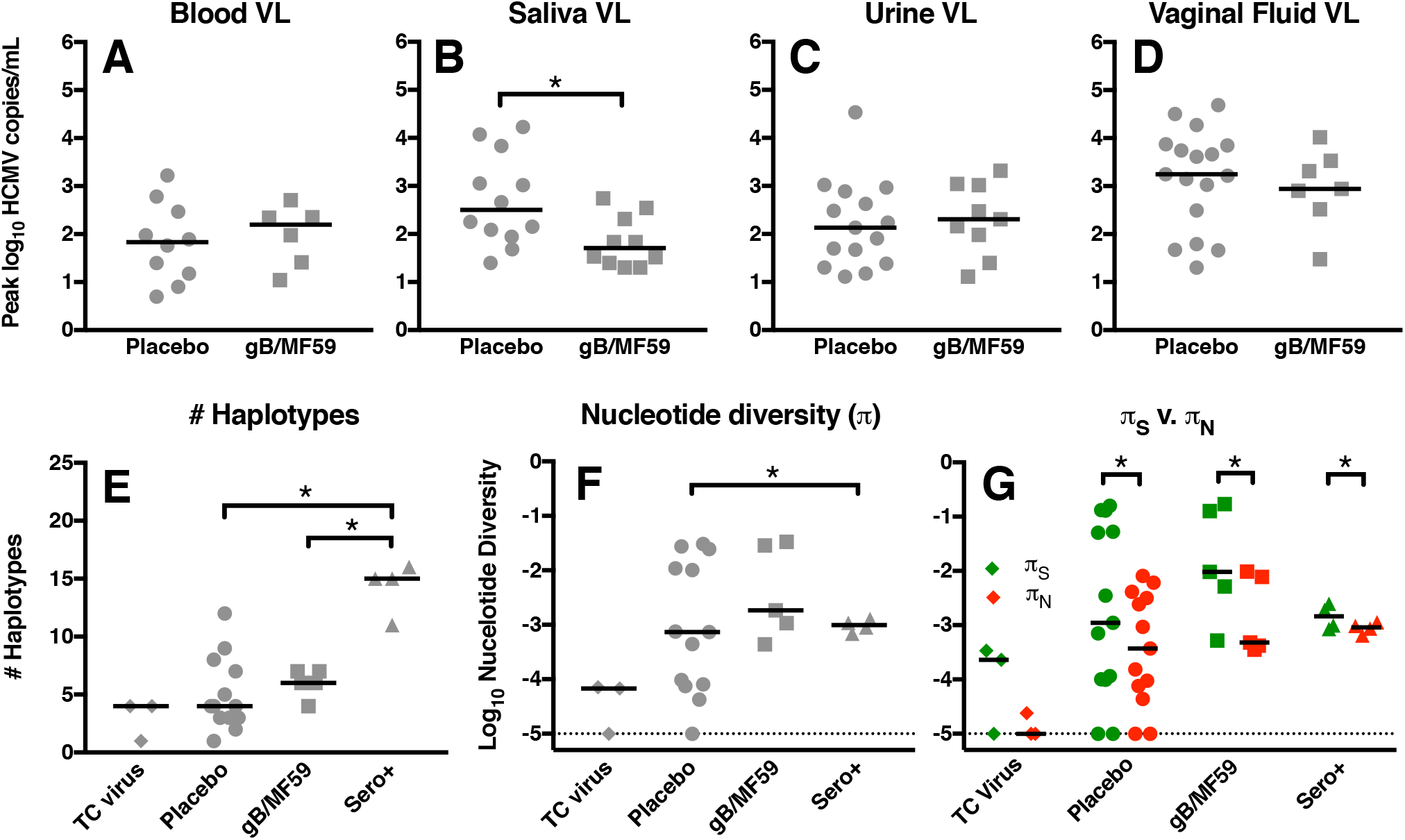
Reduced peak saliva shedding, yet similar number of viral haplotypes and nucleotide diversity between HCMV-infected glycoprotein B vaccinees and placebo recipients. Peak plasma viral load (A) as well as the peak magnitude of virus shed in saliva (B), urine (C), and vaginal fluid (D) was compared between gB vaccinees and placebo recipients. Plasma (A), urine (C), and vaginal fluid (D) viral load was not statistically different between HCMV-infected placebo recipients and gB/MF59 vaccinees, though there was reduced HCMV shedding in the saliva of vaccinees (B). The number of unique viral haplotypes (E) as well as nucleotide diversity (*π*) (F) was assessed in viral DNA amplified at the gB locus for tissue culture virus (TC virus), placebo recipients, gB/MF59 vaccinees, and seropositive, chronically HCMV-infected individuals (Sero+). (G) The magnitude of nucleotide diversity resulting in synonymous (**!**s) vs. nonsynonymous changes (!n) was compared. Horizontal bars indicate the median values for each group. *p<0.05; statistical tests employed include: viral load - Exact Wilcoxon Rank Sum test, haplotypes & *π* - Kruskal-Wallis test + post hoc Exact Wilcoxon Rank Sum test, !s vs. !n - Wilcoxon Signed Rank test.

Next, short (~550 base pair), variable regions within gB and UL130 (membrane glycoprotein targeted by neutralizing antibodies, but not included in gB/MF59 vaccine) were amplified by nested PCR then deep-sequenced (Figure S1). Unique viral haplotypes were inferred by a modified SeekDeep analysis pipeline (24) (Figure S2). gB and UL130 haplotypes were obtained from a total of 14 placebo-recipients and 6 gB/MF59 vaccinees following primary infection, as well as 4 seropositive, chronically HCMV-infected individuals. Sampling was heterogeneous, though frequently samples representing multiple anatomic compartments and/or timepoints were sequenced per subject. DNA isolated from three tissue culture virus stocks were also amplified and sequenced as a genetically-homogenous comparison. The peak number of viral haplotypes among all compartments for each patient was similar between placebo and gB/MF59 vaccinees following primary HCMV infection at both the gB (Figure 1E) and UL130 loci (Figure S3A). Interestingly, the median number of gB viral haplotypes in chronically-infected seropositive individuals was noted to be 3.75 times higher than in placebo recipients (p=0.002, Kruskal-Wallis test + post hoc Exact Wilcoxon Rank Sum test) and 2.5 times higher than in gB/MF59 vaccinees (p=0.008, Kruskal-Wallis + Exact Wilcoxon Rank Sum test). However, this trend was not identified at the UL130 locus.

The peak nucleotide diversity (p) for each patient was calculated for identified haplotypes at the gB (Figure 1F) and UL130 loci (Figure S3B). There was no statistical difference in gB p between placebo recipients and vaccinees following primary HCMV infection, though seropositive individuals had 1.3-fold higher gB p in comparison to primary HCMV-infected placebo recipients (p=0.011, Kruskal-Wallis test + post hoc Exact Wilcoxon Rank Sum test). p attributable to synonymous mutations (*π*_S_) and nonsynonymous mutations (*π*_N_) was further compared within each group at the gB (Figure 1G) and UL130 (Figure S3B) loci. Again, there was no difference in *π*_S_ or *π*_N_ between infected placebo and vaccine recipients. However, gB *π*_S_ significantly exceeded *π*_N_ for primary HCMV-infected placebos (p=0.004, Wilcoxon Signed Rank test) and gB vaccinees (p=0.001, Wilcoxon Signed Rank test) as well as for seropositive, chronically HCMV-infected individuals (p=0.016, Wilcoxon Signed Rank test), indicating that purifying selection was pervasive in the viral populations of each of these groups. Of note, the enhanced nucleotide diversity of seropositive individuals over acutely-infected placebo recipients was not identified at the UL130 locus. Furthermore, at this locus *π*_S_ was also only significantly greater than *π*_N_ in the gB/MF59 vaccinee subgroup (p=0.006, Wilcoxon Signed Rank test). Overall, these data suggest that the genetic complexity of the viral population in acutely-infected vaccinees vs placebo recipients is similar, though reduced compared to the viral population in seropositive, chronically HCMV-infected individuals.

### Viral load, number of HCMV haplotypes, and sequence diversity by anatomic compartment

Viral load measured near the time of seroconversion was assessed for blood, saliva, urine, and vaginal fluid (Table 2), as previously reported on for the placebo subset of patients (25). By bivariate analysis, after accounting for multiple samples per patient, a regression analysis of log_10_ viral load on source indicated a statistically significant difference in viral copy number by anatomic compartment (P<0.001), with vaginal virus shedding loads exceeding the viral load in all other compartments. Mean viral load in vaginal swabs was 7.14 times that of oral swabs (95% CI: 1.05-50 times, p=0.044), 14 times that of urine samples (95% CI: 33.3–100 times, p=0.001) and 25 times that of whole blood samples (95% CI: 5.5–100 times, p<0.001). Finally, we observed that the percent of samples with detectable HCMV virus decreased over time in both gB vaccinees and placebo recipients. This decrease was observed in all body fluids, although levels in whole blood were already low around the time of seroconversion (Table 3).

**Table 2.** Summary statistics of viral load in different body fluids at time of seroconversion.

| **Source** | **Total # samples** | **Log<sub>10</sub> mean VL** | **Log<sub>10</sub> median VL** | **Log<sub>10</sub> VL IQR** |
| --- | --- | --- | --- | --- |
| Oral | 20 | 2.95 | 2.12 | <2 – 2.74 |
| Urine | 24 | 3.06 | <2 | <2 – 2.56 |
| Vaginal | 19 | 3.86 | 3.25 | <2 – 3.87 |
| Whole blood | 26 | 2.21 | <2 | <2 – 2.30 |

**Table 3.** Trends in percent detectable (viral load ≥100 copies/ml) over time by source.

| **Visit\*** | **Oral**<br>% Detectable (n) | **Urine**<br>% Detectable (n) | **Vaginal**<br>% Detectable (n) | **Whole Blood**<br>% Detectable (n) |
| --- | --- | --- | --- | --- |
| 0 | 55.0 (20) | 42.7 (24) | 73.7 (19) | 26.9 (26) |
| 1 month | 75.0 ( 4) | 42.9 (7) | 83.3 (6) | 25.0 (4) |
| 2-36 months | 20.0 (20) | 23.8 (21) | 22.2 (18) | 5.6 (18) |

\*Visit 0 is around the time of seroconversion, visit 1 is one month later, etc.

The peak HCMV load for each patient in each anatomic compartment was compared (Figure 2A). Median peak vaginal HCMV shedding load was noted to be 17.95 times higher than HCMV viremia (p=0.002, Pairwise Wilcoxon Signed Rank test) and 10.72 times higher than urine HCMV shedding load (p=0.001, Pairwise Wilcoxon Signed Rank test). There were no statistical differences in the peak number of viral haplotypes identified or peak nucleotide diversity between blood, saliva, urine, or vaginal fluid at either the gB (Figure 2B,C) or UL130 (Figure S3C,D) loci. Of note, the nucleotide diversity of HCMV in whole blood was 11.54-fold higher than that of HCMV shed in urine at the gB locus and 1.8-fold higher at the UL130 locus, which is consistent with previous observations (12), though this finding was not statistically significant. Finally, we observed that *π*_S_ significantly exceeding *π*_N_ in blood (p=0.027, Wilcoxon Signed Rank test), saliva (p=0.008, Wilcoxon Signed Rank test), urine (p=0.011, Wilcoxon Signed Rank test), and vaginal fluid (p=0.010, Wilcoxon Signed Rank test) at the gB locus (Figure 2D), as well as in urine at the UL130 locus (p=0.006, Wilcoxon Signed Rank test) (Figure S3D), again indicating purifying selection in these viral populations.

**Figure 2.**
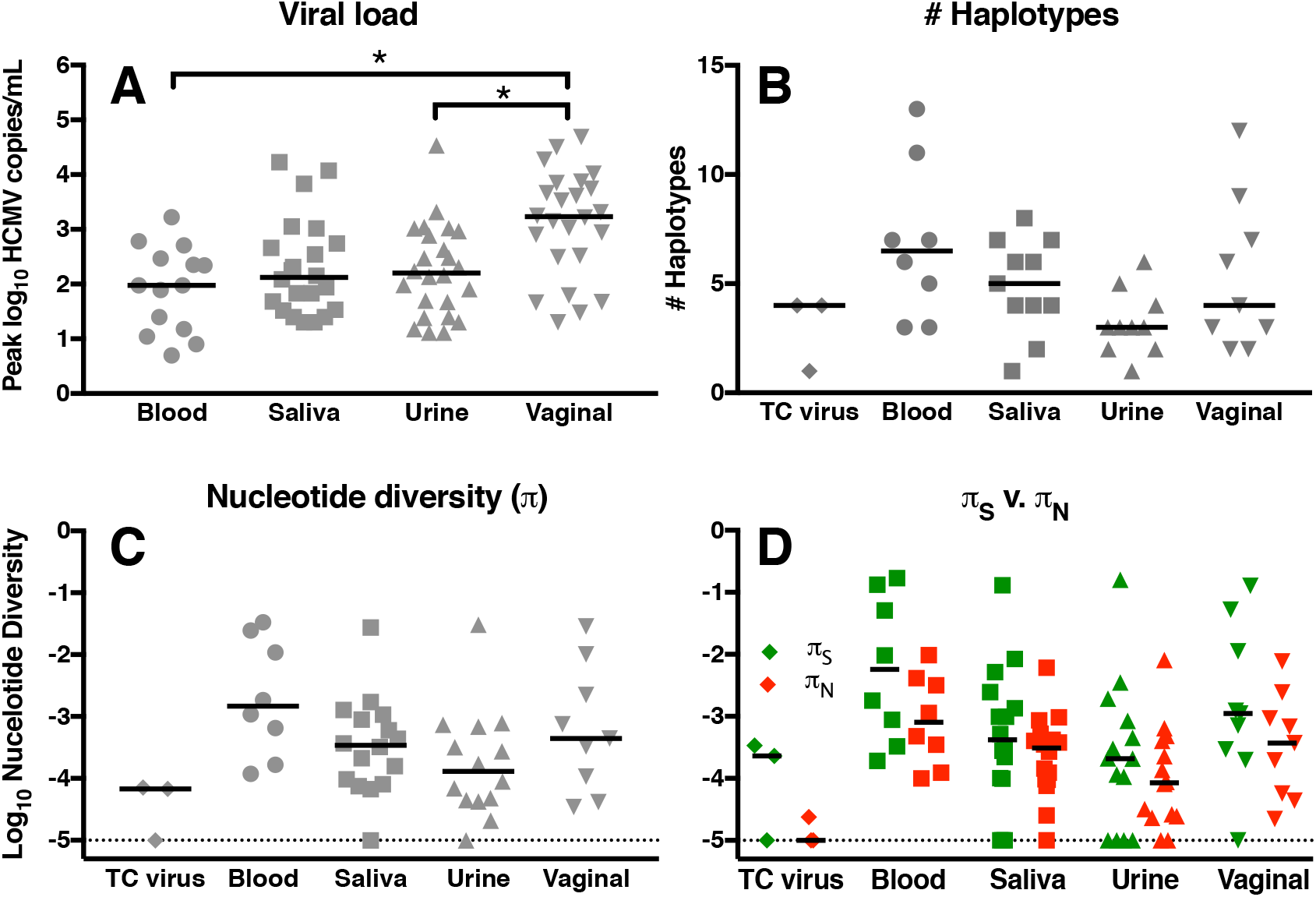
High magnitude viral shedding in vaginal fluid, yet similar number of unique viral variants and nucleotide diversity between anatomic compartments. Peak viral load was compared between anatomic compartments (A), revealing high-magnitude HCMV shedding in vaginal fluid. The number of unique viral haplotypes (B) as well as nucleotide diversity (*π*) (C) were defined for viral DNA amplified at the gB locus for tissue culture virus (TCV), as well as virus isolated from whole blood, saliva, urine, and vaginal fluid from acutely-infected gB vaccinees and placebo recipients as well as chronically HCMV-infected individuals. (G) The magnitude of nucleotide diversity resulting in synonymous (*π*_S_) vs. nonsynonymous changes (*π*_N_) was compared. Horizontal bars indicate the median values for each group. *p<0.05; statistical tests employed include: viral load - Friedman test + post hoc Pairwise Wilcoxon Signed Rank test, haplotypes & *π* - Kruskal-Wallis test + post hoc Exact Wilcoxon Rank Sum test, *π*_S_ vs. *π*_N_ - Wilcoxon Signed Rank test.

### Presence and persistence of low-frequency, unique HCMV variants

The relative frequency of unique viral haplotypes was identified for tissue culture virus stocks, chronically HCMV-infected individuals, and primary HCMV-infected gB/MF59 vaccinees and placebo recipients at the gB (Figure 3) and UL130 (Figure S4) genetic loci. For all patients and at both loci, there is typically a single dominant viral variant, with a frequency approaching 100% (range: 54–99.6%). Along with this dominant variant, we find multiple low-frequency variants, often with a frequency of less than 1% of the population, that are genetically-distinct from the dominant variant and persist over time. For example, longitudinal haplotypes identified in placebo recipients 1 and 12 indicate the persistence of both the dominant and low frequency variants from one time point to the next, suggesting that these identified variants are not simply sequencing artifact.

**Figure 3.**
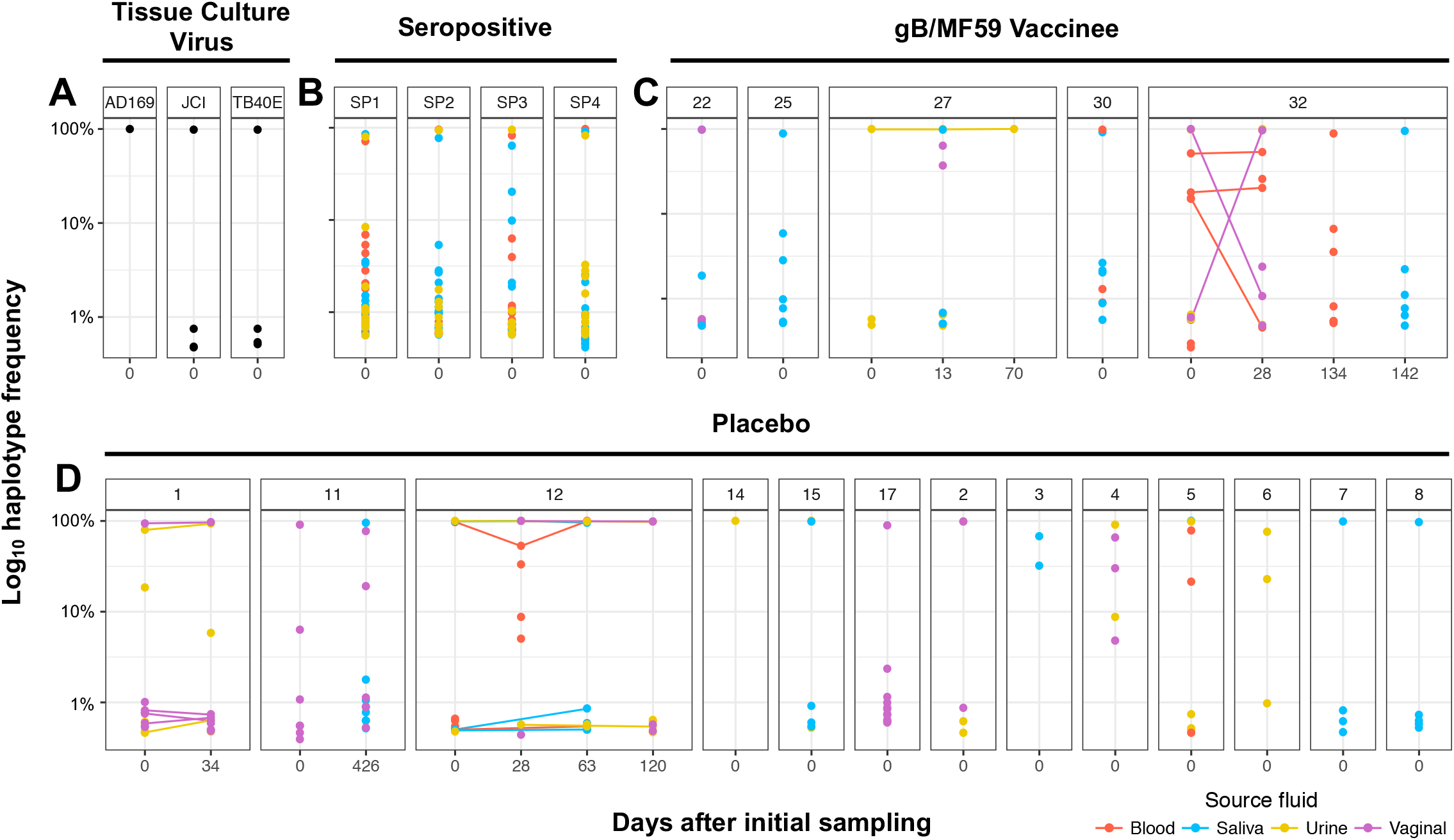
Large number of low-frequency viral variants detected at gB locus in both primary HCMV-infected and chronically-infected individuals. The relative frequency of each unique gB haplotype identified by SNAPP is displayed by individual patient and time point of sample collection. Tissue culture viruses (A) exhibited reduced population complexity by comparison. In primary HCMV-infected placebo recipients (C) and gB vaccinees (D), as well as chronically HCMV-infected women (B), there are typically one or more high-frequency haplotypes representing the dominant viral variants within the population which is accompanied by haploptyes at very low frequency representing minor viral variants (<1% of viral haplotype prevalence). Color indicates source fluid: red=blood, blue=saliva, yellow=urine, and pink=vaginal fluid. All haplotypes displayed exceed the 0.44% threshold of PCR and sequencing error established for the SNAPP method (see materials and methods for detail).

### Anatomic compartmentalization of HCMV populations in gB/MF59 vaccinees

A panel of tests for genetic compartmentalization reliant upon 6 distinct distance and tree-based methods was employed to assess the extent to which viral populations in different anatomic compartments of a single subject constitute distinct populations (26). We classified a viral population as compartmentalized when at least 3 of the 6 tests gave a statistically-significant result. Given this definition, anatomic compartmentalization at the gB locus was observed for 1 of 7 placebo recipients, 3 of 4 gB vaccine recipients, and 0 of 4 seropositive, chronically HCMV-infected individuals (Figure 4A). This frequency of genetic compartmentalization between placebo recipients and gB vaccinees approached statistical significance (p=0.088, Fisher’s Exact test), with increased compartmentalization in the vaccinee group. This same trend was not observed at the UL130 locus, as 2 of 9 placebo recipients, 1 of 4 gB vaccinees, and 2 of 3 seropositive individuals exhibited evidence of anatomic virus population compartmentalization (Figure S5). This is perhaps due to either: 1) local tissue adaptation focused at the gB but not UL130 locus or 2) lower levels of UL130 genetic diversity leading to an inability to identify compartmentalization at this locus. The pool of gB haplotypes for 3 representative individuals is shown chronologically and separated by anatomic compartment to demonstrate patients either lacking (Figure 4B) or exhibiting (Figure 4C,D) evidence of gB variant genetic compartmentalization.

**Figure 4.**
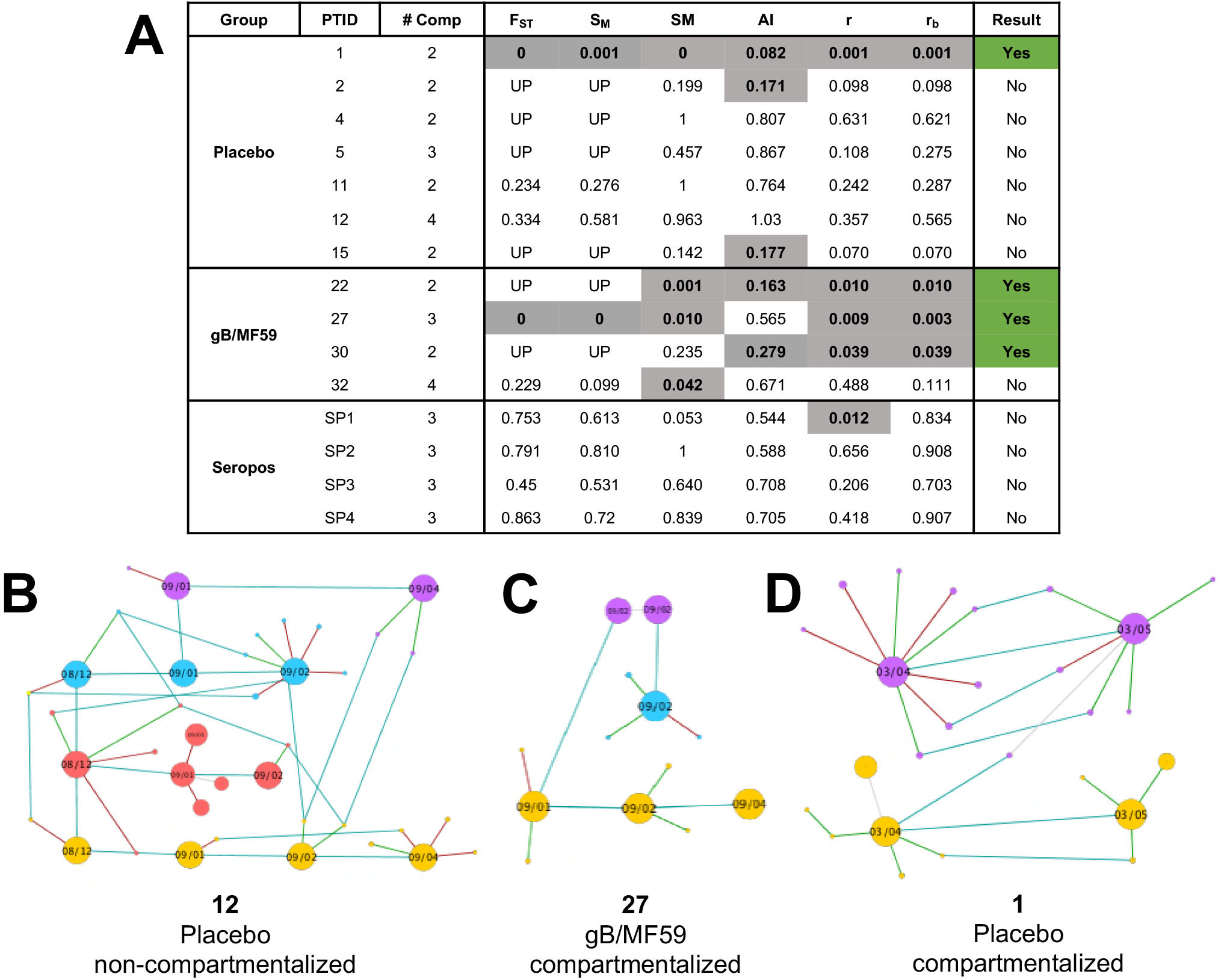
Evidence of viral genetic compartmentalization at gB locus in 3 of 4 gB vaccinees. (A) Table indicating the results of 6 distinct tests of genetic compartmentalization performed on the pool of unique gB haplotypes identified per patient, including Wright’s measure of population subdivision (F_ST_), the nearest-neighbor statistic (S_nn_), the Slatkin-Maddison test (SM), the Simmonds association index (AI), and correlation coefficients based on distance between sequences (r) or number of phylogenetic tree branches (r_b_). For each test, >1000 permutations were simulated. Significant test results suggesting genetic compartmentalization are shown in gray with bold text. Values for F_ST_, S_nn_, SM, r, and r_b_ represent uncorrected p-values, with p<0.05 considered significant. An AI<0.3 was considered a significant result. Three or more positive tests per patient was considered strong evidence for genetic compartmentalization, indicated in green. **UP** = under-powered (fewer than 5 haplotypes were present in each compartment, making F_ST_ and Snn error-prone) (B-D) Network of unique viral haplotypes by individual patient, with 1 patient lacking tissue compartmentalization (B) and 2 patients demonstrating strong evidence of viral genetic compartmentalization (D). Samples are organized chronologically from left to right, with blood shown in red, saliva in blue, urine in yellow, and vaginal fluid in purple. The size of each node reflects the relative prevalence of each haplotype. Light blue lines connect identical viral variants between time points and compartments, green lines connect variants with a synonymous mutation, and red lines those with a nonsynonymous mutation.

### gB genotype analysis and force-of-infection modeling

In addition to SNAPP, the full gB ORF was amplified, fragmented, and sequenced by NGS to identify a gB consensus sequence for each unique sample (Figure S1). Reassuringly, there was a high level of agreement of the gB genotype identified in primary HCMV-infected women between previously-published Sanger sequencing data (10), full gB ORF NGS, and SNAPP (Table 4). As previously noted by Sanger sequencing, full gB ORF NGS indicated relatively few incidences of mixed infection observed between physiologic compartments or distinct time points. However, the SNAPP technique identified additional, low-frequency viral variants corresponding to diverse gB genotypes, likely only discernable due to the enhanced sensitivity of this technique for low-frequency variants. Indeed, 11 of 18 (61%) of trial participants had evidence of minor gB genotypes by SNAPP in comparison to 2 of 32 (6%) by Sanger sequencing.

**Table 4.**
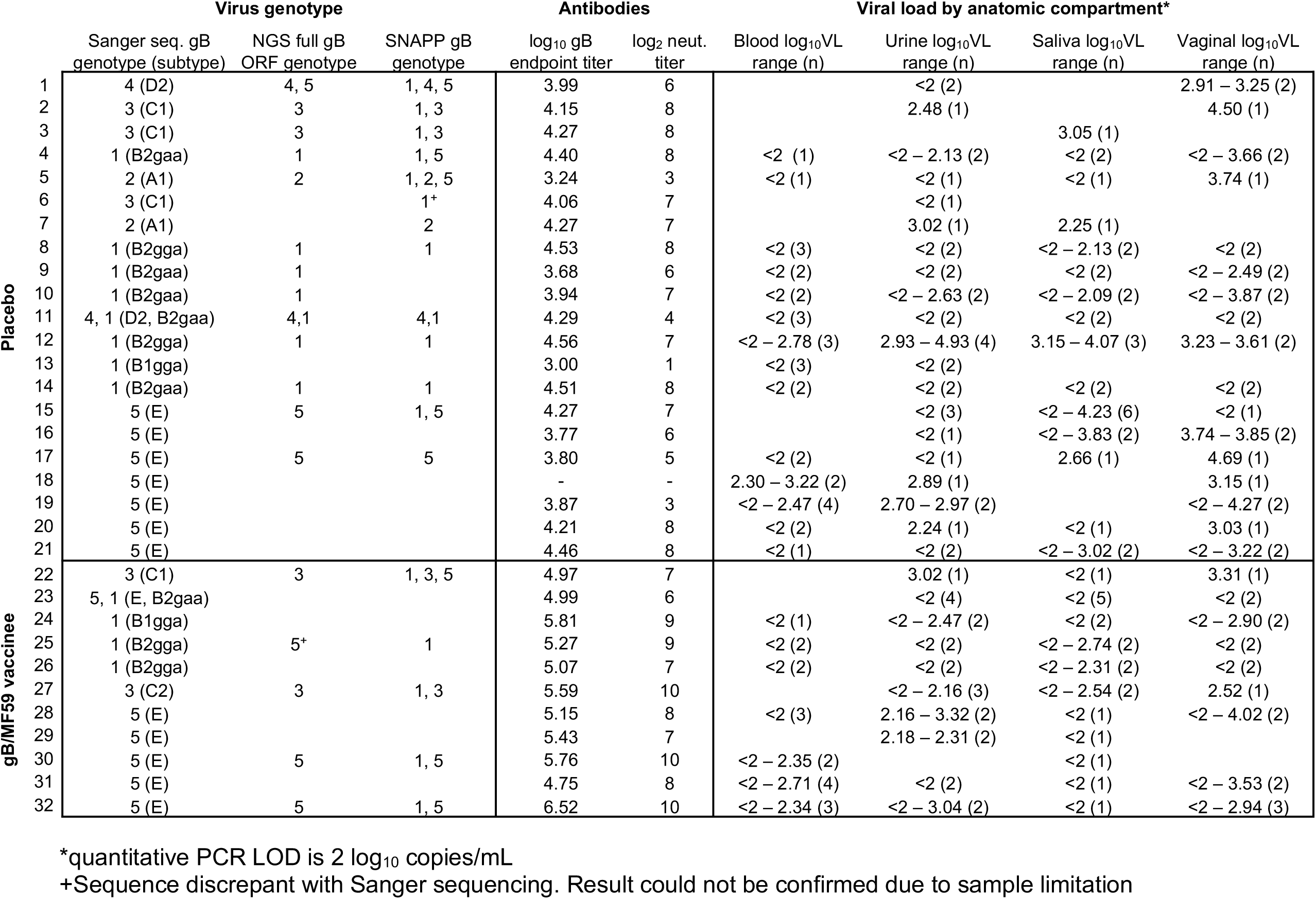
Viral load and gB genotypes detected in clinical samples from placebo recipients and gB/MF59 vaccinees using different sequencing methodologies.

We previously reported no association between gB genotype and vaccination status using Sanger sequencing data (10). However, here we observed that gB/MF59 vaccinees relatively infrequently acquired gB1, gB2, and gB4 genotype viruses. Based on phylogenetic relatedness, we hypothesize that these 3 gB genotypes could be considered to comprise a gB1/2/4 genotype “supergroup” (Figure 5A). Indeed, full gB ORF NGS data indicate that 9 of 13 placebo recipients acquired a gB1/2/4 virus compared with 0 of 5 vaccinees (Figure 5B), which was a statistically significant result (p=0.029, Fisher’s Exact test). Sanger sequencing data show a similar trend with 11 of 21 placebo recipients acquiring a gB1/2/4 virus compared with 3 of 11 vaccinees, though this difference was not significant (p=0.266, Fisher’s Exact Test).

**Figure 5.**
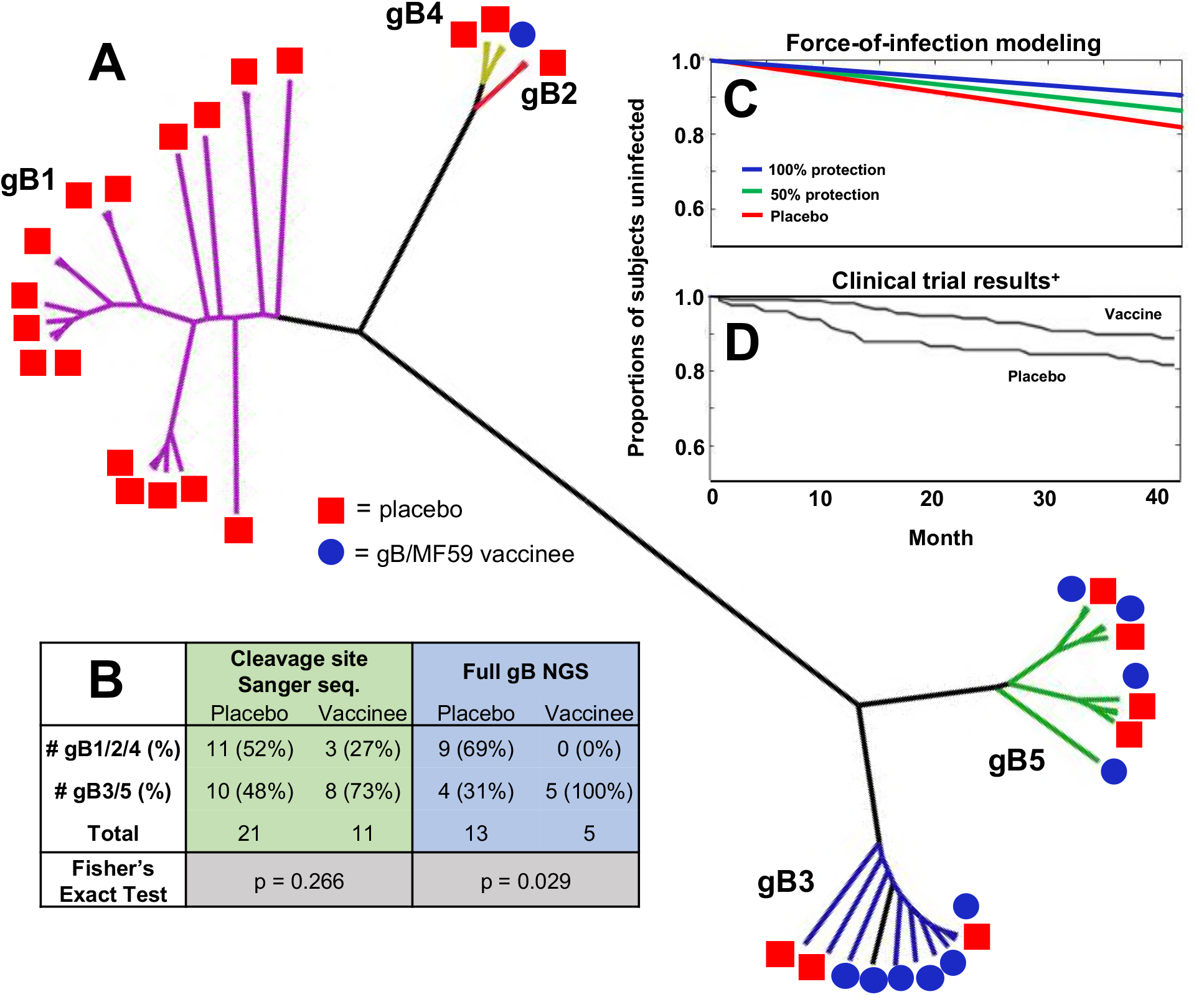
Possible protection against gB1/2/4 genotype supergroup viruses in gB/MF59 vaccinees. (A) Unrooted phylogenetic tree in a polar layout constructed using full gB open reading frame consensus sequences for each sequenced sample (often multiple samples and multiple compartments for each trial participant). Clades representing gB genotypes are highlighted in different colors: gB1=purple, gB2=yellow, gB3=blue, gB4=red, and gB5=green. (B) Number of distinct gB vaccinees and placebo recipients who acquired viral variants belonging to two supergroups of genetically-similar gB genotypes (gB1/2/4 and gB 3/5) as defined by Sanger sequencing of the cleavage site (highlighted in green) and NGS of the full gB protein (highlighted in blue). (C) Force-of-infection modeling closely predicts observed gB/MF59 vaccine trial efficacy (D) - figure adapted from Pass et al., *New England Journal of Medicine,* 2009. Force-of-infetion model iterations include that gB vaccinees are universally-protected against acquisition of gB1/2/4 variants (blue line; consistent with full gB NGS data) or 50% protected (green line; consistent with Sanger sequencing data). Model makes assumption that 1) viruses belonging to the gB1/2/4 genotype supergroup represent 52% of the circulating virus pool and 2) that the HCMV force-of-infection is 5.7 per 100 person-years.

**Figure 6.**
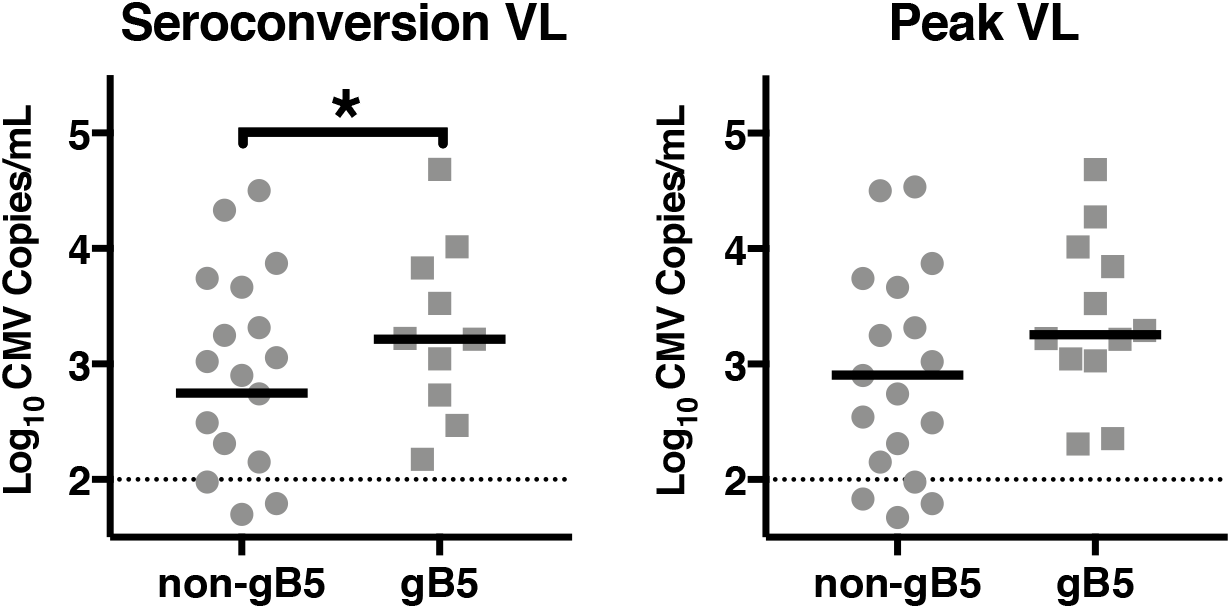
Higher viral load among women that acquired gB5 genotype viruses. A linear regression analysis of log_10_ viral load on genotype was performed at time of seroconversion (A) as well as peak viral load (B). At time of seroconversion, the viral load among women who acquired a gB5 genotype virus was 3.44 time greater than that of women shedding non-gB5 genotype virus (95% CI 1.13–10.51, p=0.031). Solid line for each grouping indicates mean value, whereas dotted black line indicates threshold of qPCR detection (100 copies/mL).

We sought to investigate whether vaccine-elicited protection against gB1/2/4 genotype viruses could have explained the partial vaccine efficacy observed in the gB/MF59 clinical trial (5) by modeling the HCMV force-of-infection (Figure 5C). Assuming that gB1/2/4 genotype supergroup comprises 52% of the circulating virus pool (11 of 21 acquired viruses among placebo recipients), we modeled the theoretical protection against gB1/2/4 variants assuming that vaccinees were either universally protected (blue line) or 50% protected (green line) in comparison to placebo recipients (red line). If we assume that vaccinees were ~50% protected against gB1/2/4 variants (reflects Sanger sequencing data - most complete data set), the modeled vaccine efficacy is 26%. Thus, force-of-infection model results are consistent with those observed in clinical trial (5), but do not fully explain the observed 50% vaccine protection.

### Viral load correlation to gB-binding antibodies, HCMV neutralization, and HCMV virus genotype

Regression analysis was performed to assess a possible correlation between viral load and the magnitude of gB-binding or HCMV-neutralizing antibodies elicited by vaccination. For both binding and neutralization, separate analyses were performed using 1) all measured viral load values (Figure S7A,B) or 2) only the highest viral load value in each participant (Figure S7C,D). No statistically significant correlation between antibody responses and viral load was identified for either the highest measured viral load (gB antibodies: R^2^=0.0428, p=0.26; neutralizing antibodies: R^2^=0.0035, p=0.75) or when all viral load measurements were included (gB antibodies: R^2^=0.0035, p=0.75; neutralizing antibodies: R^2^=0.0009, p=0.71).

Additionally, we investigated the relationship between viral load and gB genotype in 30 women using Sanger sequencing-assigned gB genotypes (most complete data-set). Two participants that did not have viral load measurments during seroconversion were excluded. A relatively high viral load was observed among participants infected with gB5 genotype viruses in both vaccine and placebo recipients. To investigate further, we dichotomized genotype as gB5 versus all other gB genotypes, then performed a linear regression of log_10_ viral load on genotype. Around the time of seroconversion, a statistically significant association was found between log_10_ viral load and genotype, with the mean viral load in women with gB5 genotype viruses being 3.44 times greater than that of women shedding non-gB5 genotype viruses (95% CI 1.13–10.51, p=0.031). When considering all visits, this association was no longer statistically significant; however, this is likely a result of the decrease in viral load over time observed in all HCMV-infected participants.

## Discussion

Despite the partial efficacy demonstrated by gB/MF59 vaccination in multiple clinical trials (5–7), there has been little examination of the impact of this vaccine on the *in vivo* viral populations. In this investigation, we sought to employ the enhanced sensitivity of next-generation sequencing (NGS) technology to delve deeper into the question of whether there are discernable differences between viruses acquired by gB/MF59 vaccinees and placebo recipients. The advantage of NGS over more traditional sequencing methodologies is the ability to detect minor viral variants, which contribute to the diversity of the overall viral population (Figure 3, Figure S4). We discovered that numerous minor viral haplotypes, exceeding the threshold of PCR and sequencing error, were detectable in nearly all clinical samples tested, which is consistent with results of HCMV whole-genome sequencing that have suggested numerous genetic variants at <1 *%* frequency in the viral population (11–13). Interestingly, seropositive women reliably had more gB haplotypes (Figure 1E) than acutely-infected vaccinated subjects, indicating a higher number of genetically-distinct viral variants in seropositive, chronically HCMV-infected individuals. This observation complements previous work demonstrating that recently-seroconverted young women have very low incidence of mixed infection, yet that multiple gB genotypes are almost universally detectable in chronically-HCMV infected individuals (22). Altogether, these data favor a model that mixed infections in healthy individuals result from independent, sequential infection events.

Throughout the study, we identified several indications of vaccine-mediated effects on the viral population. First, peak HCMV shedding in saliva was reduced by an order of magnitude in gB/MF59 vaccinees compared to placebo, though no difference was observed for peak systemic viral load or peak viral shedding in urine and vaginal fluid. This finding suggests that gB-elicited antibodies may have limited viral replication in salivary glands. gB/MF59 vaccination is known to elicit high titers of gB-specific IgG, IgA, and SIgA in parotid saliva (27), which may have suppressed HCMV salivary replication and reduced saliva viral shedding. Furthermore, we observed a trend towards reduced viral load in vaccinees when considering samples from all timepoints or those solely from the timepoint of seroconversion, though these comparisons were not statistically significant. Analogously, serum antibody gB-binding and HCMV-neutralization did not correlate with viral load at peak or at the time of seroconversion. These data suggest altogether that gB/MF59 vaccination had minimal quantifiable impact on *in vivo* HCMV replication. Indeed, our observation that gB5 genotype viruses were associated with increased viral load further suggests that host response may have a lesser impact on viral replication during acute infection than strain identity and viral pathogenicity.

The second identified impact of vaccination on the viral population was anatomic compartmentalization. We found that 3 of 4 gB vaccinees with viral DNA available from multiple compartments exhibited evidence of viral genetic compartmentalization at the gB locus, in contrast to only 1 of 7 placebo recipients. As has been previously described (10), we observed that the dominant viral haplotype was identical between anatomic compartments in the majority of subjects. The evaluation of gB-specific compartmentalization was therefore only possible because of the ability of Short NGS Amplicon Population Profiling (SNAPP) to detect minor viral variants in clinical samples. Our data are consistent with HCMV whole-genome NGS indicating tissue-specific variants with intrahost variable SNPs present at relatively low frequency (11, 12). The mechanism leading to compartmentalization in vaccinees is unclear, though it is possible this phenomenon stems from either neutral or positive selection in distinct anatomic compartments. One hypothesis is that systemic gB-specific antibodies restricted free dissemination of HCMV variants between tissue compartments. Such a bottleneck might have reduced founder population size and increased the speed and likelihood of stochastic fixation of neutral SNPs and formation of genetically-distinct viral populations (28, 29). Alternatively, it is possible that local factors including cell type and local antibody production/secretion at the site of HCMV replication selected for “more fit” viral variants in each compartment.

Lastly, we observed an interesting trend that gB/MF59 vaccine recipients had reduced acquisition of gB1/2/4 genotype supergroup viruses. This finding was statistically significant when considering the genotypes identified by full gB ORF NGS, though not significant when considering a larger dataset of the genotypes previously defined by Sanger sequencing (Figure 5, Table 4) (10). Of note, the gB immunogen in this vaccine trial was based on the Towne strain (gB1 genotype prototypic sequence) suggesting the possibility of vaccine-mediated protection against viruses with genetic/antigenic similarity to the vaccine strain. Complementarily, we have observed in the same gB/MF59 vaccinee cohort that gB-elicited neutralization activity was increased against the autologous Towne strain virus, but not heterologous viruses AD169 and TB40/E (8). These observations raise the possibility that gB/MF59 vaccine-elicited protection against HCMV acquisition had a limited antigenic scope. As demonstrated by force-of-infection modeling, gB1/2/4 supergroup-specific protection could partially explain the 50% vaccine efficacy observed in this phase 2 clinical trial. The concept of neutralization breadth has not been explored extensively for HCMV, though several studies have described strain-specific neutralization (30-32). However, it is important to place this finding in context. Low frequency gB1/2/4 haplotypes were detectable in additional vaccinees by SNAPP (Table 1), suggesting there may not have been a true barrier to gB1/2/4 *acquisition* but perhaps merely restricted gB1/2/4 virus replication. Furthermore, since the sample size is not robust, it is possible that our observation may have been the result of sampling bias.

Far and away the largest limitation of this study was sample availability. Unfortunately, the scope of our investigation was restricted by: 1) the original sampling timeline employed during the clinical trial, 2) the availability of clinical samples, and 3) the integrity of the DNA more than a decade following DNA extraction. Additionally, as with any study based on PCR amplification and DNA sequencing, there is a potential for primer bias, contamination, and background error to obscure the results. We instituted several measures to increase data integrity. First, primers were designed and validated to prevent amplification bias (23). Additionally, PCR, sequencing, and analysis was completed in duplicate to reduce the likelihood of contamination affecting results. An advantage of this investigation is that we were able to validate our two sequencing methodologies (SNAPP and full gB ORF NGS) by comparing observed gB genotypes with previously published data (10). The gB genotype predicted by Sanger sequencing and NGS sequencing methodologies were identical for 88% of all samples. However, because of the relatively small cohort size and potential for sequencing error, our observed trends certainly merit further investigation.

Nonetheless, this investigation is the first to employ NGS of viral DNA from infected HCMV vaccinee and placebo recipients in an attempt to characterize the viral determinants of HCMV acquisition. Importantly, our data robustly support notions in the HCMV field that the *in vivo* viral population has a level of genetic complexity that cannot be appreciated using conventional sequencing methodologies (11–13), though the biological relevance of this genetic diversity remains to be determined. Furthermore, our observation of reduced saliva shedding and a high rate of gB sequence compartmentalization in vaccinees suggests a definable impact of gB-elicited antibodies on viral population dynamics. Finally, the possible vaccine-elicited protection against gB1/2/4 supergroup viruses observed in this study is intriguing, implying that that antigen-specific immune responses may have played a role in vaccine protection. These discoveries are hypothesis-generating for future explorations into the mechanism anti-HCMV antibody-mediated immunity and ought to be investigated further in a larger cohort.

## Materials and Methods

### Study population

The study population was comprised of 32 postpartum women who acquired HCMV infection while participating in a phase 2, randomized, double-blind, placebo controlled clinical trial of an HCMV vaccine (10). 21 women received placebo, while 11 received the gB/MF59 vaccine. Clinical trial participants were HCMV-seronegative, healthy postpartum women immunized with gB protein (Sanofi Pasteur; based on Towne strain, gB1 genotype) + MF59 squalene adjuvant (Novartis) or placebo. Subjects were vaccinated on a 0, 1 and 6 month schedule and were screened for HCMV infection every three months for 3.5 years using an antibody assay which detects seroconversion to HCMV proteins other than gB (33). Subjects with serologic evidence of infection were tested for HCMV in blood, urine, saliva and vaginal swab from one month to 3.5 years after seroconversion. Samples were collected from 30 women around the time of acute seroconversion (21 placebo and 9 vaccinee). Across all patients and time points, a total of 201 samples (oral n=50, urine n=58, vaginal n=45, and whole blood n=48) were collected during visits ranging from 1 to 36 months. The total number of samples per patient ranged from 1 to 12 (median=7). While a maximum of 7 oral samples was available per patient, the maximum number of urine, vaginal and whole blood samples was 4, 3, and 4, respectively. Aliquots of each specimen were stored at −80°C. Institutional review board (IRB) approval was obtained from University of Alabama at Birmingham and Johns Hopkins Hospital and all subjects signed an approved consent form. The Duke University Health System determined that analysis of de-identified samples from these cohorts does not constitute human subjects research.

### DNA extraction and quantitative PCR

HCMV DNA was extracted from 400 μL of original samples - blood, urine, saliva, or vaginal swab in culture media - using the MagAttract virus mini M48 kit (Qiagen) on the BioRobot M48 instrument. The quantitative PCR assay is based on amplification of a 151-bp region from the US17 gene (25, 34). As previously reported, the limit of detection is 100 copies/mL (4 copies/reaction), with a measurable range of 2.0 to 8.0 log_10_ copies per mL.

### Antibody assays

IgG to HCMV gB was measured using an ELISA method. The vaccine antigen, a recombinant HCMV gB molecule (Sanofi Pasteur; based on Towne strain, gB1 genotype), was used to coat 96-well plates. Eight serial dilutions starting at 1:200 were tested. A horseradish peroxidase (HRP)-conjugated goat anti-human IgG (1:50,000 in HRP stabilizer, KPL) was used as detection antibody, followed by development using TMB peroxidase substrate solution (TMB 2-Component Microwell Peroxidase Substrate Kit, KPL). Endpoints were calculated based on the intersection between the straight line through the falling portion of the optical density-serum dilution curve and the cut-off optical density below which sera were negative for antibodies to gB. Neutralizing antibodies were measured using a plaque reduction assay with HCMV AD169 (ATCC #CCL-171) and MRC-5 cells (ATCC #VR-538) in 24-well tissue culture plates. Duplicate two-fold dilutions of sera were tested starting at 1:8. Plaque counts were performed after a 7-day incubation and the highest serum dilution with ≥ 50% reduction in plaque count was considered the endpoint titer.

### Short NGS amplicon population profiling (SNAPP)

Flow chart detailing the sequencing strategy is shown in Figure S1. Variable regions approximately 550 base-pairs in length within gB (UL55) and UL130 were amplified in duplicate by a nested PCR using the primers denoted in Table S1. Overhang regions were conjugated to PCR2 primers for subsequent Illumina index primer addition and sequencing: forward primer overhang = 5’-TCGTCGGCAGCGTCAGATGTGTATAAGAGACAG-[locus]-3’ and reverse primer overhang = 5’-TCTCGTGGGCTCGGAGATGTGTATAAGAGACAG-[locus]-3’. Template DNA extracted from primary fluids was added to 50 μl of 1x PCR mixture containing 100 nM of each primer, 2 mM MgCl_2_, 200 μM each of dNTP mix (Qiagen), and 0.2 U/μl Platinum Taq polymerase. PCR reactions consisted of an initial 2-minute denaturation at 98°C, followed by 35 PCR cycles (98°C for 10 seconds, 65°C for 30 seconds, and 72°C for 30 seconds), and a final 72°C extension for 10 minutes. Following each amplification step, products were purified using Agencourt AMPure XP beads (Beckman Coulter). Illumina Nextera XT index primers were added by 15 cycles of amplification. The indexed PCR product was run on a 1 % agarose gel, then gel-purified using ZR-96 Zymoclean Gel DNA Recovery Kit (Zymogen). The molar amount of each sample was normalized by real-time PCR using the KAPA library amplification kit (KAPA Biosystems). The library of individual amplicons was pooled together, diluted to an end concentration of 14 pM, combined with 20% V3 PhiX (Illumina), and then sequenced on Illumina Miseq using a 600-cycle V3 cartridge (Illumina).

### Full glycoprotein B open reading frame PCR and sequencing

Flow chart denoting sequencing strategy is shown in Figure S1. The full gB open reading frame (ORF) was amplified by nested PCR using the primers denoted in Table S1. Template DNA extracted from primary fluids was added to 50 μl of 1x PCR mixture containing 100 nM of each primer, 2 mM MgCh, 200 μM each of dNTP mix (Qiagen), and 0.2 U/μl Platinum Taq polymerase. PCR reactions consisted of an initial 2-minute denaturation at 98°C, followed by 35 PCR cycles (98°C for 10 seconds, 65°C for 30 seconds, and 72°C for 3 minutes), and a final 72°C extension for 10 minutes. Following each amplification step, products were purified using Agencourt AMPure XP beads (Beckman Coulter). The PCR2 product was run on a 1% agarose gel, then gel extracted using the ZR-96 Zymoclean Gel DNA Recovery Kit (Zymogen). Purified product was tagmented using the Nextera XT library prep kit (Illumina). Subsequently, Nextera XT index primers were added to the tagmented DNA by 15 cycles of amplification. The molar amount of each sample was normalized by real-time PCR using the KAPA library amplification kit (KAPA Biosystems). The library of individual amplicons was pooled together, diluted to an end concentration of 14 pM, combined with 20% V3 PhiX (Illumina), and then sequenced on Illumina Miseq using a 600-cycle V3 cartridge (Illumina).

### SNAPP haplotype reconstruction and nucleotide diversity

Data processing flow chart is shown in Figure S2. First, raw paired-end reads were merged using the PEAR software under default parameters (35). The fused reads were then filtered using the extractor tool from the SeekDeep pipeline (http://baileylab.umassmed.edu/SeekDeep) (24), filtering sequences according to their length, overall quality scores, and presence of primer sequences. All filtered sequencing reads were included for subsequent haplotype reconstruction using the qluster tool from SeekDeep. This software accounts for possible sequencing errors by collapsing fragments with mismatches at low-quality positions. For each given sample, haplotypes had to be present in both of 2 sample replicates to be confirmed. On average, concordance between the replicates was quite high as assessed by linear regression correlation and slope of the relative frequencies of each haplotype (Figure S6). Each gB haplotype was assigned to 1 of 5 described gB genotypes by assessing the shortest genetic distance (nucleotide substitutions) between the haplotype and reference gB genotype sequences (36). Nucleotide diversity (*π*) was computed as the average distance between each possible pair of sequences (37):

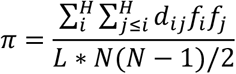

Where *L* = sequence length in nucleotides for *π*, *N* = Total number of reads in sample, *dij* = Number of nucleotide differences between haplotype i and *j, fi* = Number of reads belonging to haplotype *i*. *π*_N_*S* and *π*_S_ were calculated as the average *dS* and *dN* between pairs of haplotypes weighted by the haplotypes abundance:

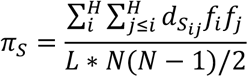

Where *L* = sequence length in amino acids for *πN, π*_S_*, N* = Total number of reads in sample, *dSij* = *dS* between haplotype *i* and *j* sequences, *fi* = Number of reads belonging to haplotype *i*. Correlations were performed between various measures of viral population diversity (viral load, number of haplotypes, *π*, *π*_S_, and *π*i_n_), and suggest that haplotypes, *π*, *π*_S_, and *π*i_n_ are somewhat related measures although are not directly equivalent (Figure S8).

### Assessment of anatomic compartmentalization of virus populations

A panel of tests using diverse analytical methods is hypothesized to be the most accurate means to infer tissue compartmentalization (26). Thus, we selected six tests employing both distance-based and tree-based algorithms. Wright’s measure of population subdivision (F_ST_, distance-based) compares mean pairwise genetic distance between sequences from the same compartment to that of sequences from the same compartment (38). The nearest-neighbor statistic (Snn, distance-based) measures how frequently the nearest neighbor to each sequence is in the same or different compartments (39). The Slatkin-Maddison test (SM, tree-based) calculates the minimum number of migration events between compartments, compared to the number of migration events expected in a randomly-distributed population (40). The Simmonds association index (AI, tree-based) examines the complexity of the phylogenetic tree structure (41). Finally, correlation-coefficients (r and r_b_, tree-based) correlates distances between sequences in a phylogenetic tree with compartment of origin based either on distance between sequences (r) or number of tree branches between sequences (r_b_). Distance-based tests used the TN93 distance matrix (42), and tree-based methods employed a neighbor-joining algorithm. Distance-based tests were not conducted for patients with fewer than 5 haplotypes per compartment since this is known to produce unreliable results (26). All tests were conducted using HyPhy software (https://veg.github.io/hyphy-site) (version 2.22), with test statistics estimated from 100+ permutations. For each test, a p-value of <0.05 or an association index <0.3 was considered statistically significant. Three or more positive test results from these six test statistics was considered strong evidence for compartmentalization.

### Phylogenetic trees and genotype assignment

Protein or nucleotide sequences of interest were aligned using the ClustalW algorithm (43) in Mega (version 6.06) (44). A neighbor-joining tree was constructed using the Los Alamos National Labs “neighbor treemaker” (https://www.hiv.lanl.gov/components/sequence/HIV/treemaker/treemaker.html), then the tree was plotted in FigTree (version 1.4.3). Full gB ORF sequences were assigned to gB genotypes based on phylogenetic proximity to reference gB sequences (36). Because of sample limitation, if the full gB assigned genotype did not match the genotype assigned to SNAPP amplicons and/or previously published Sanger sequencing data (10), these sequences were omitted from the phylogenetic analysis.

### Force-of-infection modeling

The cohort of women in the postpartum gB/MF59 vaccine trial were predominantly African-American (>70%) (5), and thus we utilized an HCMV force-of-infection estimate for non-Hispanic, African-American individuals of 5.7 per 100 persons (45). Additionally, we made the assumption that gB1/2/4 genotype supergroup viruses comprise 52% of the circulating virus pool, based on 11 of 21 placebo recipients acquiring gB1/2/4 viruses. Finally, we hypothesized that gB1-vaccinated individuals were universally protected against all circulating gB1/2/4 supergroup genotype viruses. Modeling was done using Matlab, and source code is included in the supplementary material.

### Statistical analysis

Descriptive displays and statistics were used to examine the patterns in viral load measurements by vaccination status (vaccine or placebo), source of measurement (type of body fluid) and gB genotype. Although there were multiple visits and measurements for study participants over time, the analyses of viral load distribution was initially restricted to samples collected at the first clinic visit following HCMV seroconversion. Analyses were performed to compare viral load in vaccine vs placebo recipients, to determine whether viral copy number was different between body fluids, and whether specific genotypes could be associated with higher viral load. A log_10_ transformation of viral load measurement was performed, and a value of 1 was added to each zero measurement (nondetectable) prior to transformation. Simple linear regression of log_10_ viral load was performed to assess the bivariate relationship between HCMV copy number and fluid type or genotype, respectively. The regression analysis accounted for the multiple fluid samples obtained from the same patient at the same visit by clustering and calculating robust standard errors to correlate observations within a single patient. To assess trends in viral load over time, a second analysis was performed using samples collected from all visits in all participants with measurements starting from the time of seroconversion. This descriptive analysis was performed to assess the kinetics of viral load over time in the different body fluids. Data were analyzed using STATA v10.

Additionally, in a separate analysis, the peak viral load was defined for each patient and zero-value measurements excluded. Comparisons of peak viral load data between placebo and vaccine group were performed using Exact Wilcoxon rank sum test. Comparisons of peak viral load between tissue compartments were performed using Friedman test followed by pairwise Wilcoxon signed-rank test. Comparisons of peak haplotypes number and peak nucleotide diversity data between groups were performed using Kruskal-Wallis test followed by Exact Wilcoxon rank sum test. Comparisons of peak haplotypes and peak nucleotide diversity data between tissue compartments were performed using pairwise Wilcoxon signed-rank test. Comparisons of PiS and PiN within the same tissue compartment were performed using Wilcoxon signed-rank test. The associations among different assays were calculated using Kendall’s tau. The Benjamini-Hochberg FDR p-value correction was used to correct for multiple comparisons (46). A p-value less than 0.05 (2-tailed) was considered significant for all analyses. All statistical tests were completed using SAS v9.4.

### Data and materials availability

SNAPP and full gB ORF sequence data used in this manuscript is currently being compiled and will be made available in the NCBI SRA database prior to publication. Code for force-of-infection modeling will be included in supplementary material. Source data for analyses is available upon request.

## Author Contributions

C.S.N., R.A.B., and S.R.P. designed research; C.S.N., M.S., and M.F. performed research; C.S.N., D.V.C, K.K., M.D.W., and R.A.B. analyzed data; R.F.P, K.K., and R.A.B. contributed samples and expertise; and C.S.N., R.A.B., and S.R.P. wrote the paper.

## Acknowledgements

The authors recognize the Duke Viral Genetic Analysis core for assistance with sample preparation and sequencing. Additionally, we thank Nicholas Hathaway for development of the SeekDeep analysis pipeline employed for haplotype analysis. The gB/MF59 vaccine clinical trial was supported by grants to the University of Alabama at Birmingham from NIH/NIAID (P01AI043681 and U01AI063565) and by Sanofi Pasteur. This work was supported by: NIH/NICHD Director’s New Innovator grant to S.R.P (DP2HD075699), NIH/NIAID R21 to S.R.P. (R21AI136556), NIH/NIAID R08 to R.B. (R08AI074907), and NIH/NICHD F30 grant to C.S.N (F30HD089577). The funders had no role in study design, data collection and interpretation, decision to publish, or the preparation of this manuscript. The content is solely the responsibility of the authors and does not necessarily represent the official views of the National Institutes of Health.

## Supplementary Tables and Figures

**Table S1.**
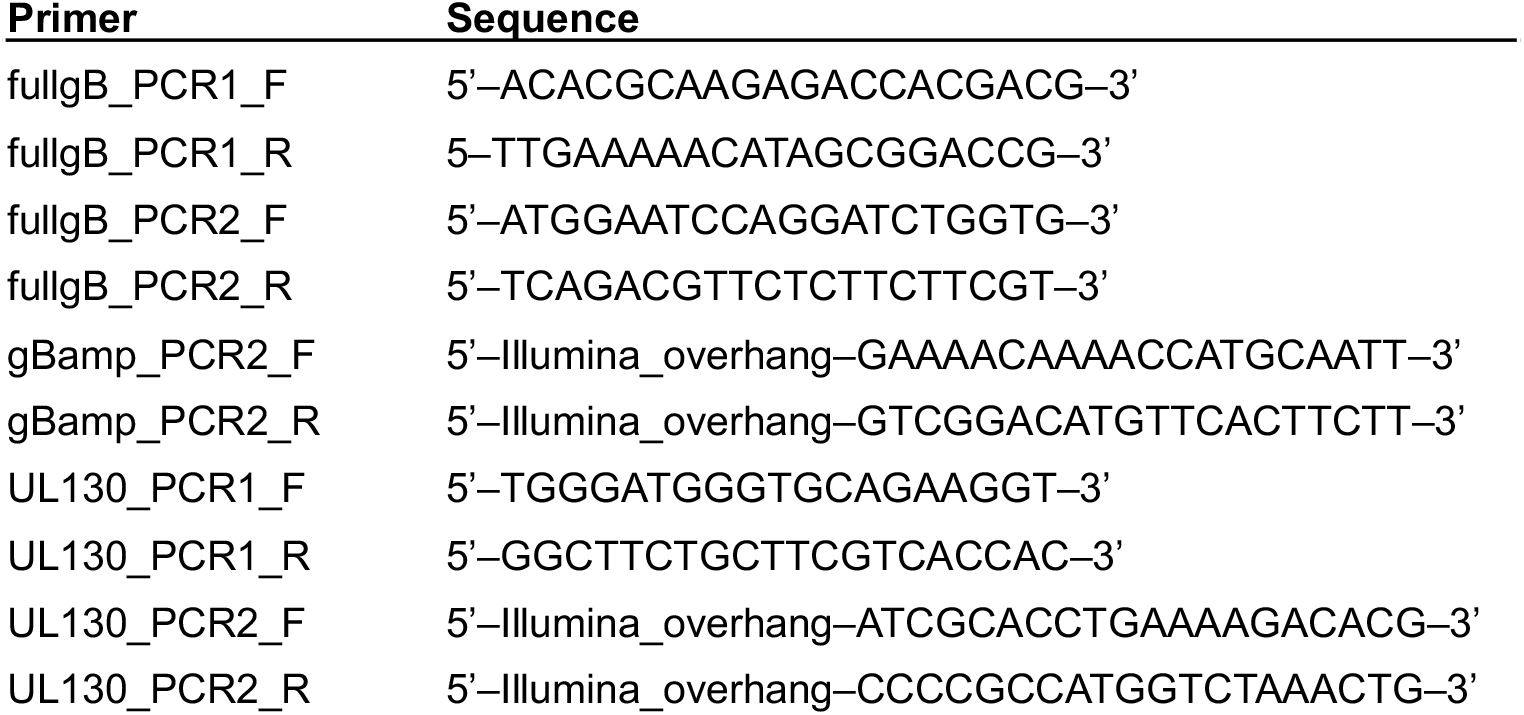
Primer sequences.

**Figure S1.**
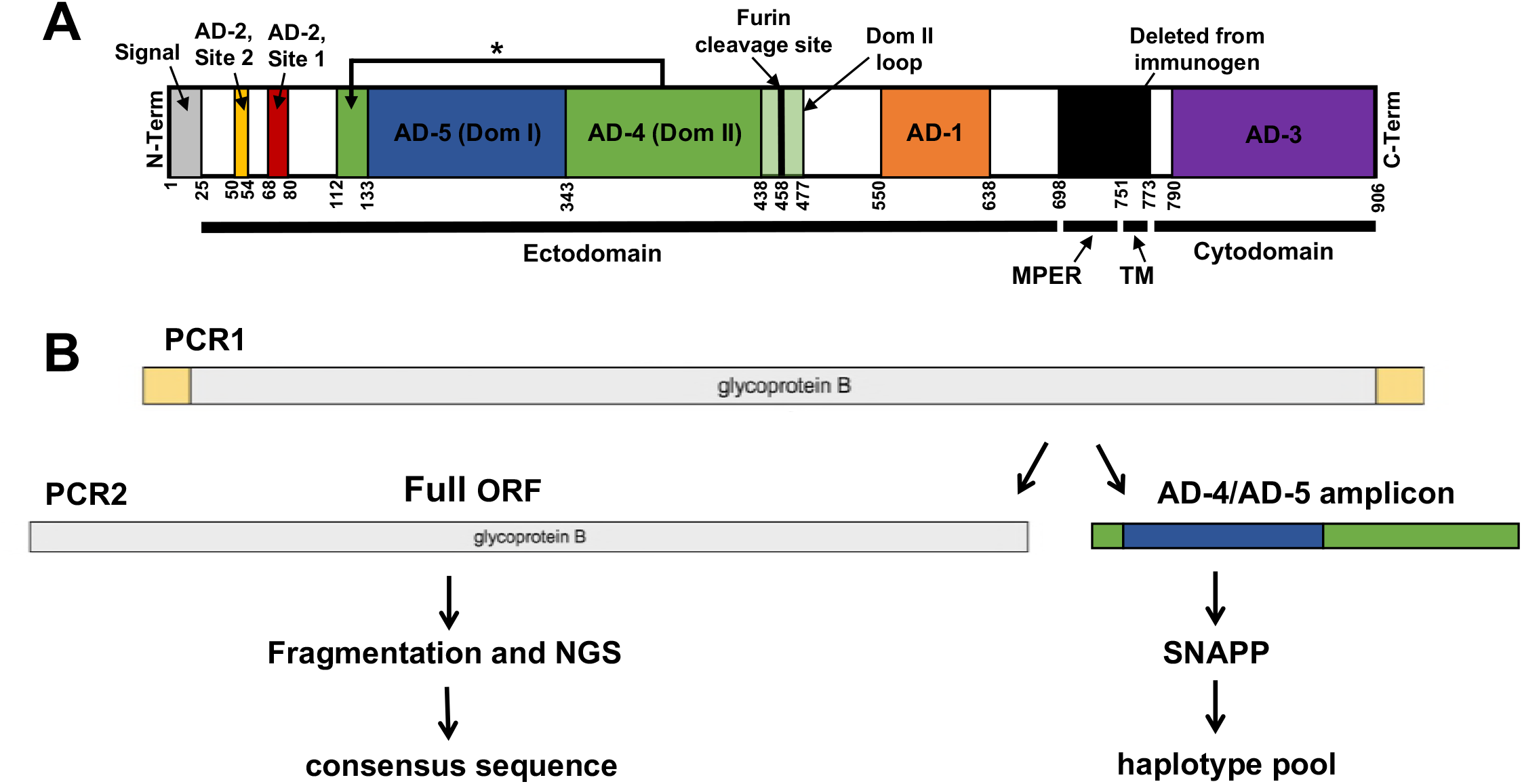
Linear structure of gB and PCR amplification/next-generation sequencing strategy. (A) The full gB HCMV open reading frame (ORF) is shown, from N-terminus on the left to C-terminus on the right. The four distinct regions of the gB structure are indicated by black bars at the base of the figure, including the ectodomain, membrane proximal external region (MPER), transmembrane domain (TM), and the cytodomain. Major antigenic regions indicated include AD-1 (orange), AD-2 site 1 (red), AD-2 site 2 (yellow), AD-3 (purple), AD-4 (Domain II) (green), and AD-5 (Domain I) (blue). Numbers indicate approximate amino acid residues dividing each region of interest. The gB immunogen employed in this clinical trial contained the full gB ORF with the furin cleavage site mutated and excluding a region from amino acid residue 698 to 773 (containing MPER and TM regions) to facilitate protein secretion during production. Diagram was adapted from Burke et al., *Plos Pathogens,* 2015 and Hebner et al., *Nature Communications,* 2015. (B) PCR amplification strategy consists of an initial PCR1 step with primers external to the gB ORF, followed by PCR2 amplification of the full gB ORF or an amplicon containing AD-4 and AD-5. Full gB ORF was NGS sequenced to generate a consensus sequence, while gB amplicons were sequenced directly and raw reads used to infer unique viral haplotypes.

**Figure S2.**
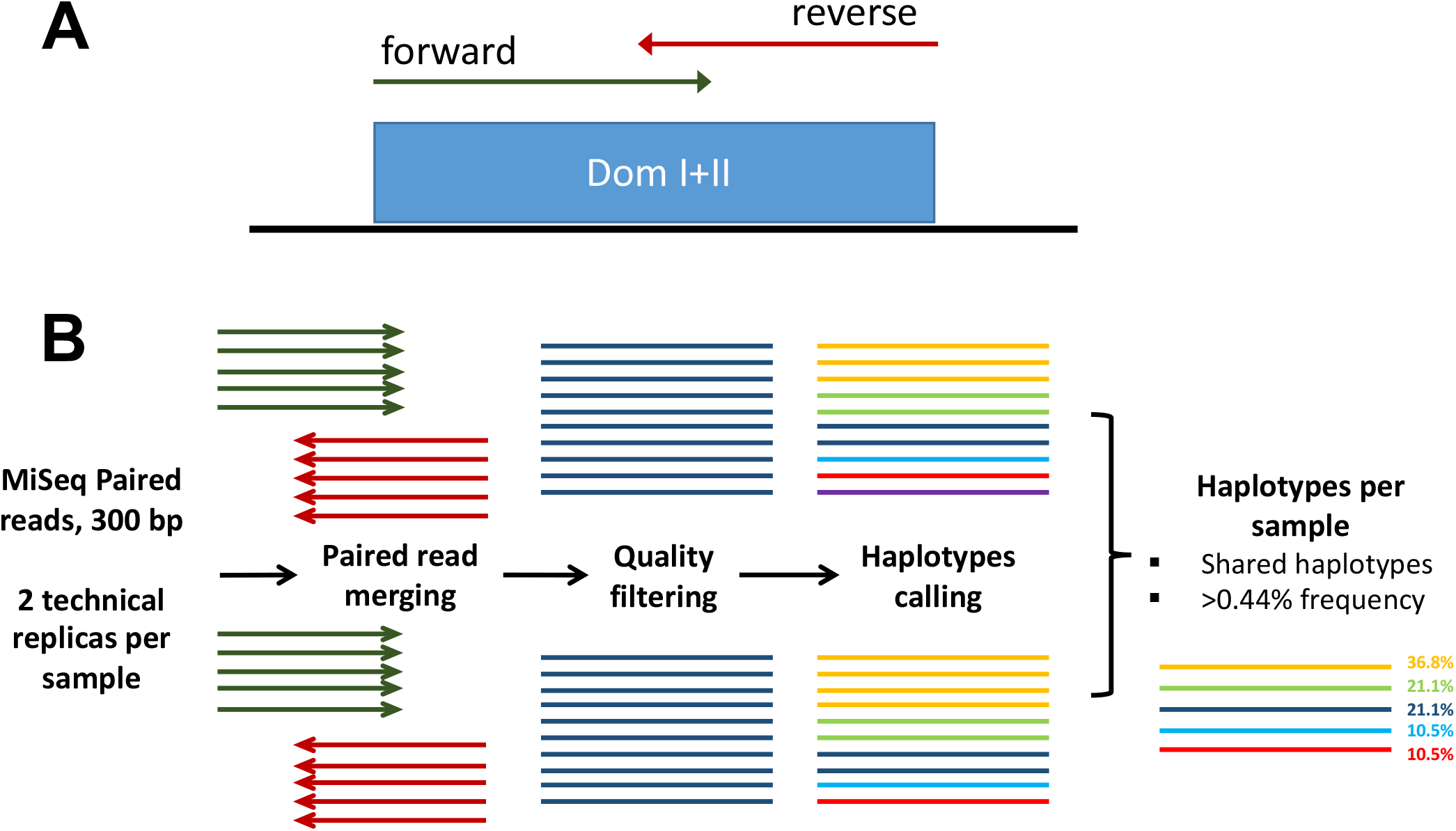
SNAPP analysis pipeline using SeekDeep. (A) Paired-end reads (forward=green, reverse=red) were obtained for an approximately 550 base-pair amplicon on an Illumina Miseq platform. (B) Paired-end reads were merged, filtered for read quality, then clustered into unique haplotypes. Haplotypes identified in both technical replicates at a frequency above the determined 0.44% cutoff were included for subsequent analysis.

**Figure S3.**
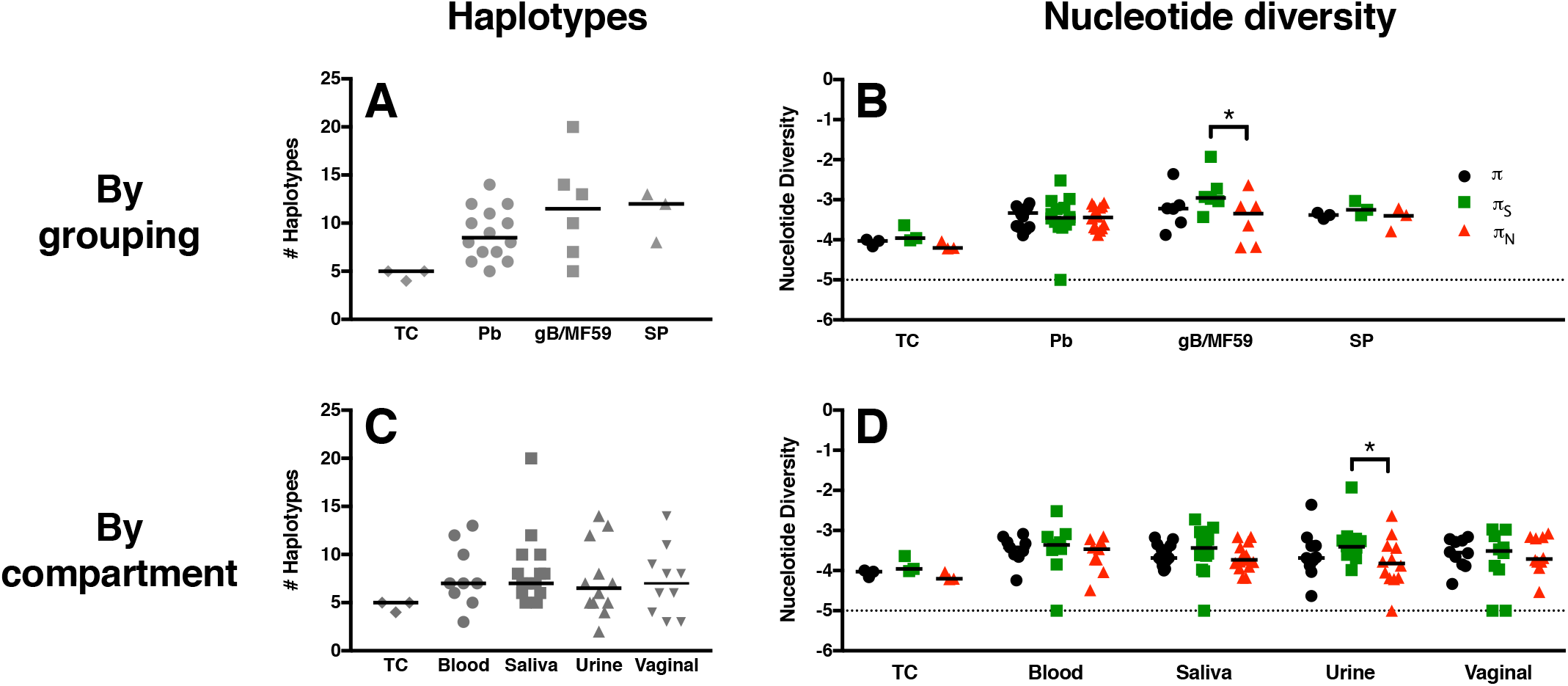
UL130 unique viral variants and peak nucleotide diversity is similar both between gB vaccine and placebo groups and between anatomic compartments. The number of unique viral haplotypes as well as nucleotide diversity (*π*) were assessed for viral DNA amplified at the UL130 locus between treatment groups (A,B) as well as between physiologic compartments (C,D). Tissue culture virus (TC virus), as well as virus isolated from whole blood, saliva, urine, and vaginal fluid. (G) The magnitude of nucleotide diversity resulting in synonymous (*π*_S_) vs. nonsynonymous (*π*_N_) changes was compared. Horizontal bars indicate the median values for each group. *p<0.05, viral load = Friedman test + post hoc Pairwise Wilcoxon Signed Rank test, haplotypes & *π* = Kruskal-Wallis test + post hoc Exact Wilcoxon Rank Sum test, *π*_S_ vs. *π*_N_ = Wilcoxon Signed Rank test.

**Figure S4.**
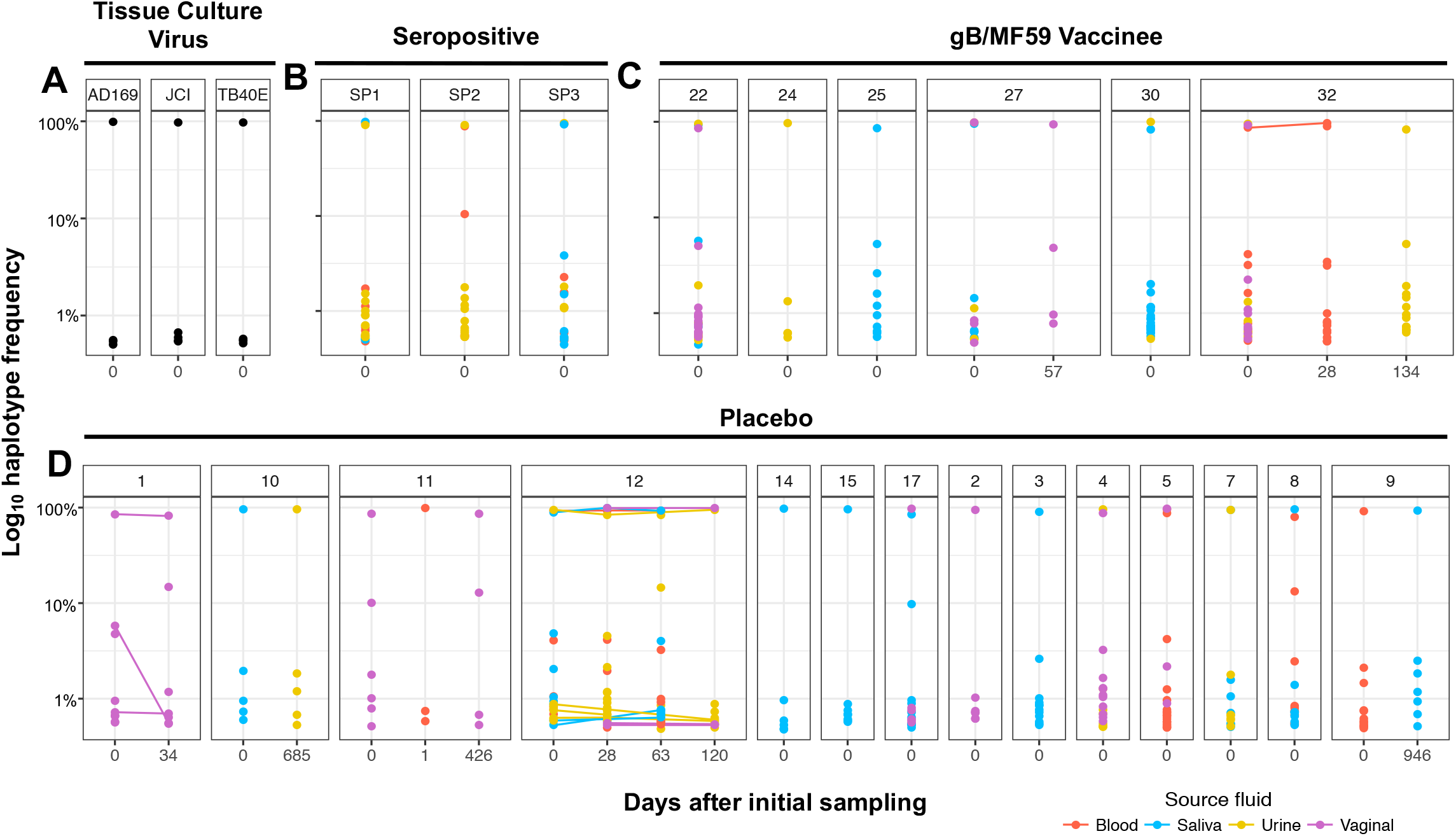
Low-frequency viral variants detectable at UL130 locus in both primary HCMV-infected and chronically-infected individuals. The relative frequency of each unique UL130 haplotype identified by SNAPP is displayed by individual patient and time point of sample collection. In primary HCMV-infected placebo recipients (A) and gB vaccinees (B) as well as chronically HCMV-infected women (C), there are typically one or more high-frequency haplotypes representing the dominant viral variants comprising the population accompanied by haploptyes at very low frequency representing minor viral variants (<1% of viral haplotype prevalence). Tissue culture viruses (D) exhibited reduced population complexity by comparison. Color indicates source fluid: red=blood, blue=saliva, yellow=urine, and pink=vaginal fluid. All haplotypes displayed exceed the 0.44% threshold of PCR and sequencing error established for the SNAPP method (see materials and methods for detail).

**Figure S5.**
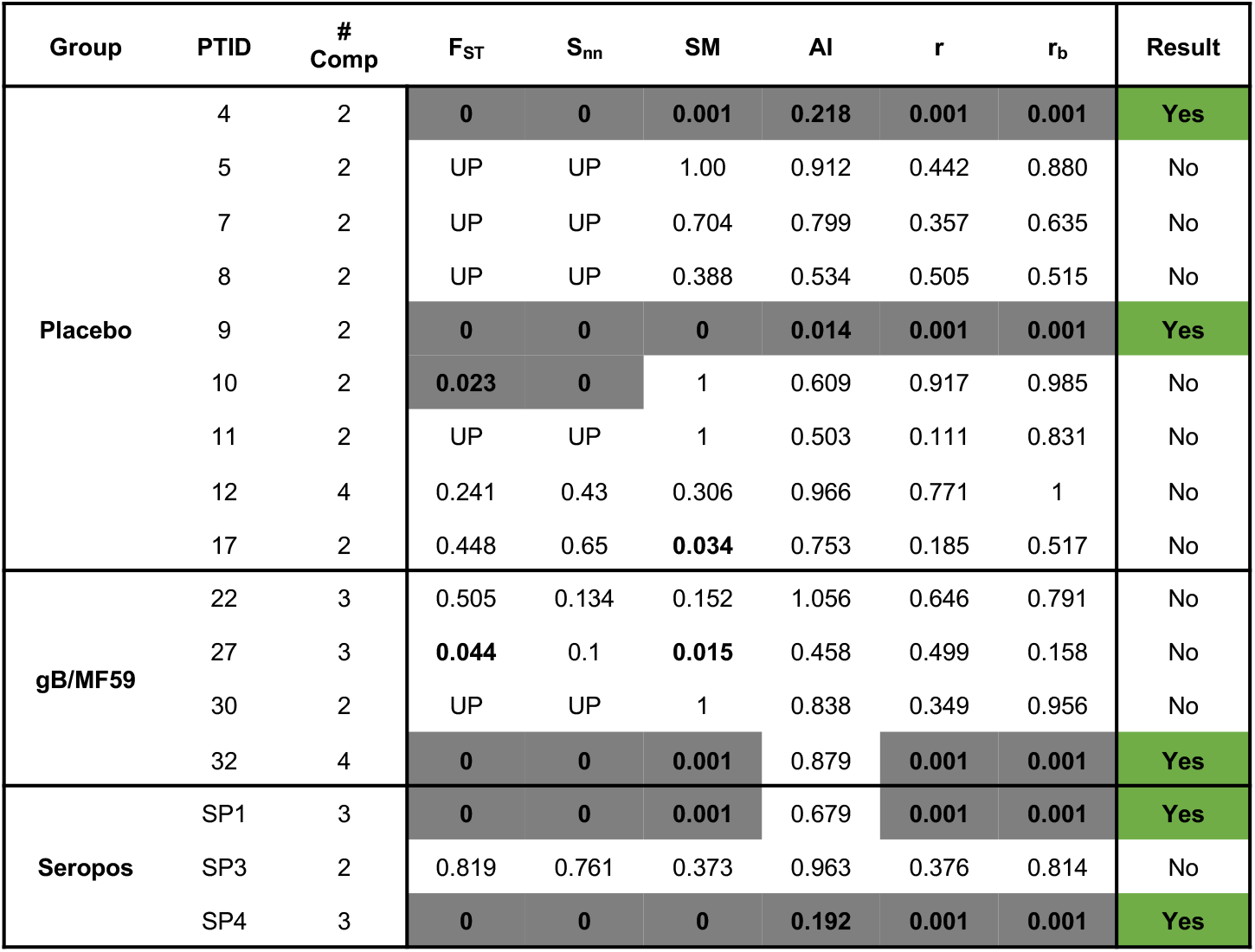
Lack of genetic compartmentalization of anatomic compartment-specific viral variants detected at UL130 locus in vaccinees. Table indicating the results of 6 distinct tests for genetic compartmentalization performed on the pool of unique UL130 haplotypes identified per patient, including Wright’s measure of population subdivision (F_ST_), the nearest-neighbor statistic (S_nn_), the Slatkin-Maddison test (SM), the Simmonds association index (AI), and correlation coefficients based on distance between sequences (r) or number of phylogenetic tree branches (r_b_). For each test, >1000 permutations were simulated. Significant test results suggesting genetic compartmentalization are shown in gray with bold text. Values for F_ST_, S_nn_, SM, r, and r_b_ represent uncorrected p-values, with p<0.05 considered significant. An AI<0.3 was considered a significant result. Three or more positive tests per patient was considered strong evidence for genetic compartmentalization, indicated in green.

**Figure S6.**
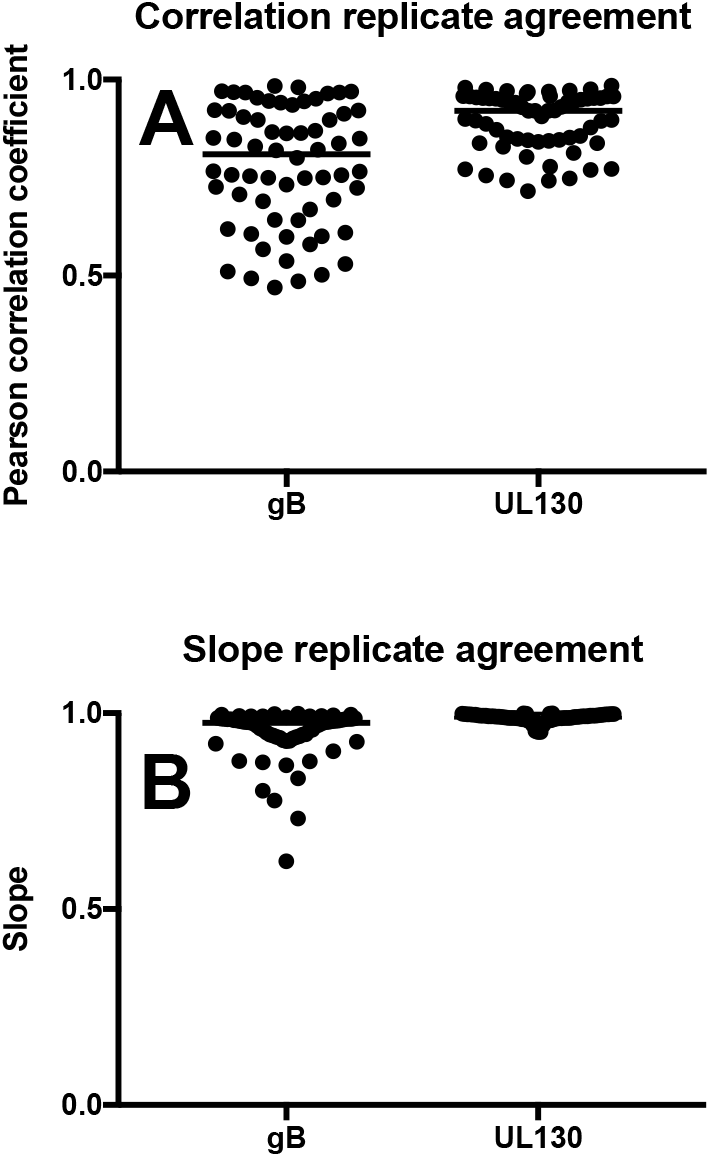
High degree of concordance in haplotype identity and frequency between sequencing replicates. Haplotype identity and frequency were calculated for two technical replicates. The correlation (A) and slope (B) of the haplotype frequencies was compared between technical replicates for both gB and UL130 amplicons, and indicate a high degree of agreement between replicates.

**Figure S7.**
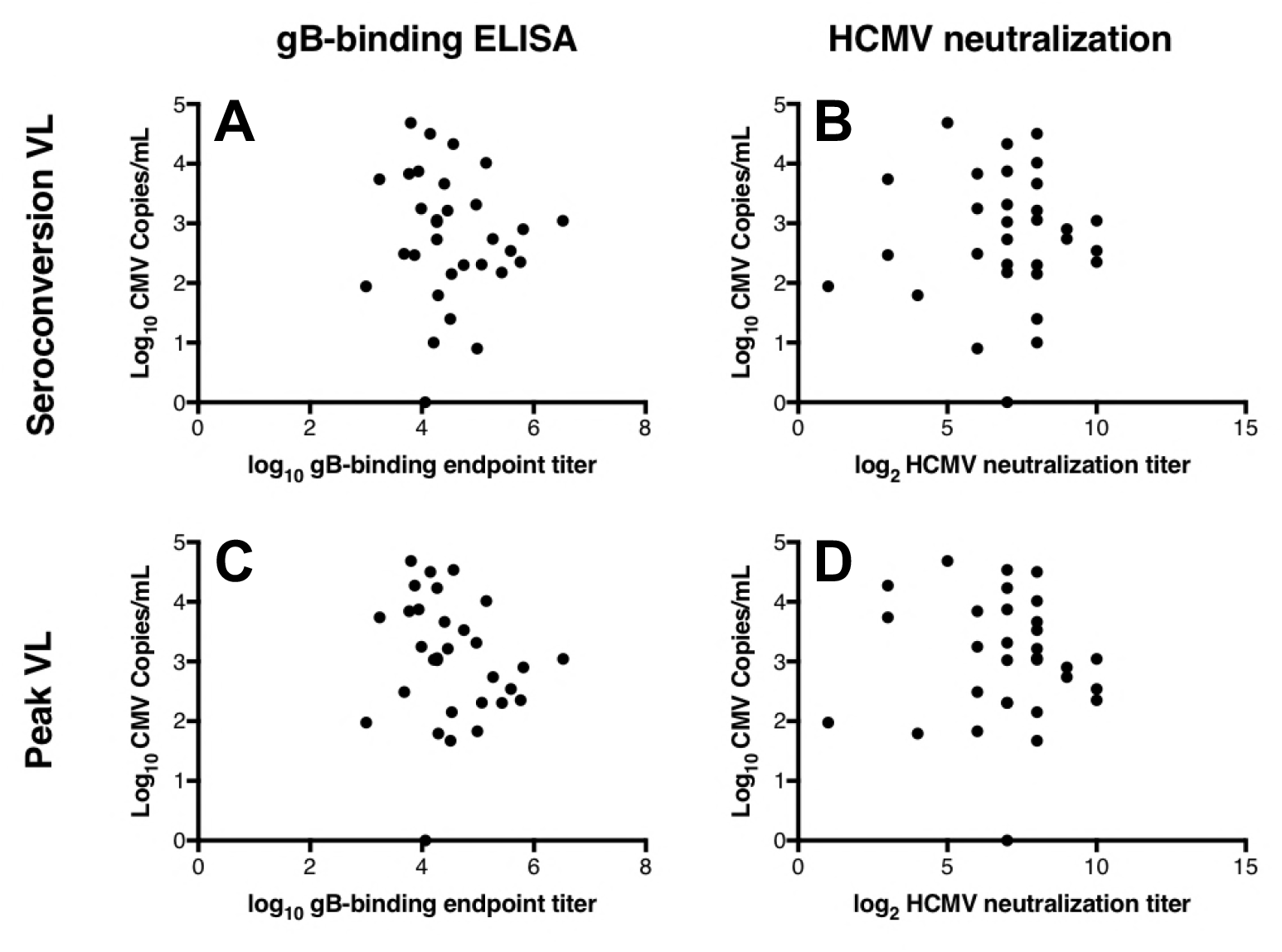
Viral load is not correlated with gB antibody-binding or HCMV neutralization. Viral load at seroconversion does not correlate with the magnitude of gB-binding (R^2^=0.0428) (A) nor with HCMV-neutralization titer (R^2^=0.0035) (B). Furthermore, peak viral load neither correlates with gB-binding (R^2^=0.0035) (C) nor neutralization titer (R^2^=0.0009) (D).

**Figure S8.**
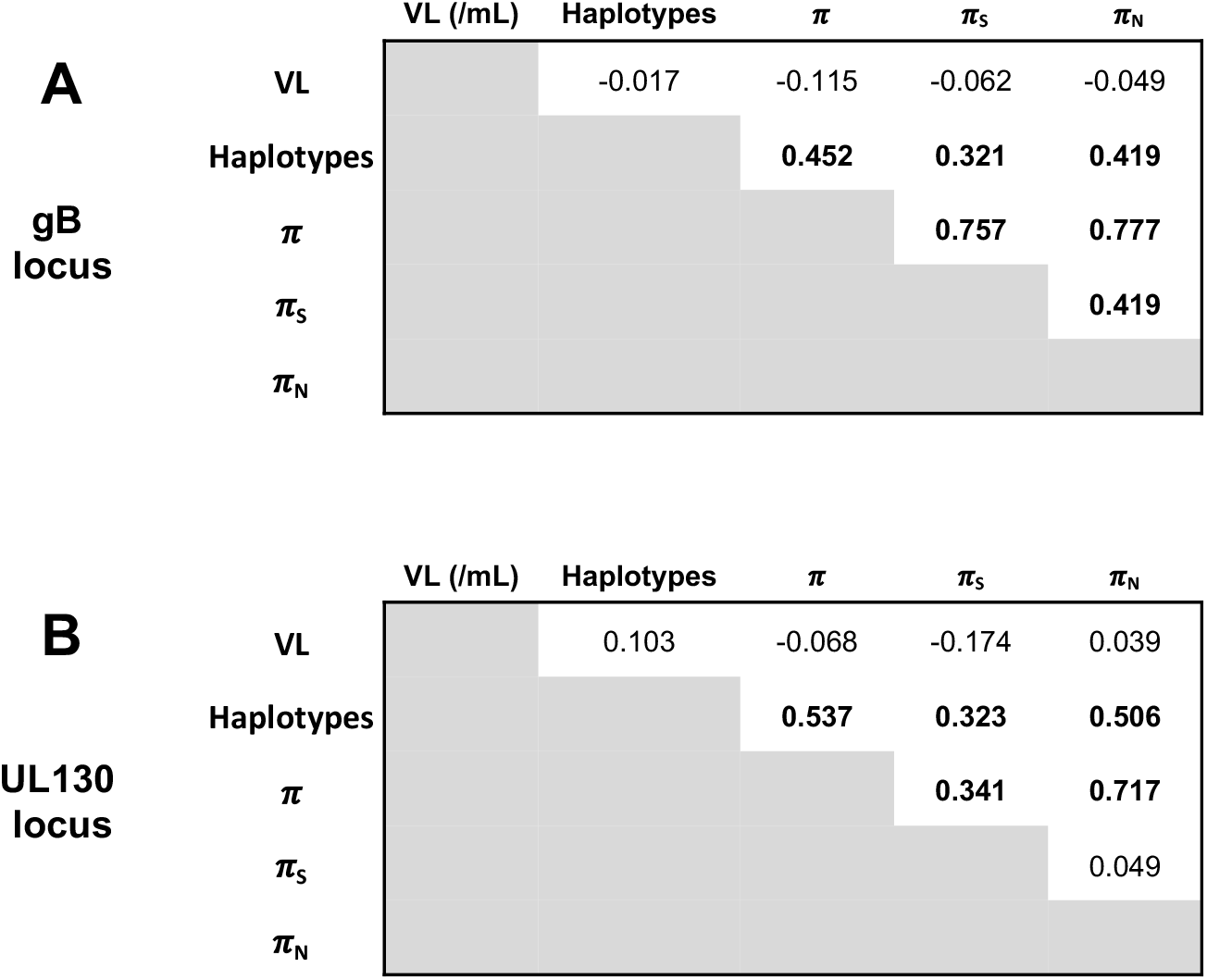
Correlations between viral load, number of unique variants, and nucleotide diversity. Kendall Tau correlation coefficients are shown for viral load (VL), number of haplotypes, nucleotide diversity (*π*), as well as synonymous nucleotide diversity (*π*_S_), and nonsynonymous nucleotide diversity (*π*_N_). Bold values indicate a significant correlation (uncorrected p<0.05)<ENT FONT=(normal text) VALUE=46>.「/ENT>

